# Enhancing tumor-infiltrating T cells with an exclusive fuel source

**DOI:** 10.1101/2024.05.20.595053

**Authors:** Matthew L. Miller, Timothy J. Thauland, Rina Nagarajan, Wenqi Ellen Zuo, Miguel A. Moreno Lastre, Manish J. Butte

## Abstract

Solid tumors harbor immunosuppressive microenvironments that inhibit tumor-infiltrating lymphocytes (TILs) through the voracious consumption of glucose. We sought to restore TIL function by providing them with an exclusive fuel source. The glucose disaccharide cellobiose, which is the building block of cellulose, contains a β-1,4-glycosidic bond that that animals (or their tumors) cannot hydrolyze, but microbes have evolved enzymes to catabolize cellobiose into useful glucose. We equipped mouse T cells and human CAR-T cells with two proteins enabling import and hydrolysis of cellobiose and demonstrated that cellobiose supplementation during glucose withdrawal restores key anti-tumor T-cell functions: viability, proliferation, cytokine production, and cytotoxic killing. Engineered T cells offered cellobiose suppress murine tumor growth and prolong survival. Offering exclusive access to a natural disaccharide is a new tool that augments cancer immunotherapies. This approach could be used to answer questions about glucose metabolism across many cell types, biological processes, and diseases.

## Introduction

Glucose is a critical fuel for cellular bioenergetics and a major source of biosynthetic precursors for anabolic pathways. Upon antigen stimulation, CD8+ T cells extensively rewire their metabolism and robustly engage aerobic glycolysis to support the various functions required of cytotoxic T cells, including survival, proliferation, cytokine production, and cytotoxicity^1–6^. For over 60 years^7^ it has been known that abnormally low glucose concentrations are found in the tumor microenvironment (TME) of solid tumors due to the voracious nature of tumor metabolism, leading to a competition for glucose that contributes to cancer progression due to stunted T-cell effector functions^8,9^. We hypothesized that TILs could be invigorated if provisioned with an *exclusive* source of glucose that was inaccessible to cancer cells, allowing them to fulfill their role in clearing tumors more effectively.

Cellobiose, a glucose disaccharide that comprises cellulose and is found abundantly in plant matter, has great potential to serve as a carbon and energy source but remains inert to catabolic processes in animal cells for two primary reasons. First, metazoan sugar transport is restricted to monosaccharides. Second, the β-1,4-glycosidic bond that joins glucose molecules in cellobiose cannot be efficiently hydrolyzed by mammalian glycoside hydrolases^10,11^. The transport and hydrolyzation of cellobiose, on the other hand, can be efficiently carried out in cellulolytic microbes such as fungi and bacteria^12–14^. Thus, cellobiose could offer an exclusive source of glucose for engineered T cells, as it is inaccessible to tumors.

Here, we report that the heterologous expression of a cellobiose transporter and a β-glucosidase from the mold *Neurospora crassa* in primary mouse and human T cells endows them with the ability to robustly catabolize cellobiose, which rescues T cells from glucose deprivation. We demonstrate that tumor cells lack the ability to use cellobiose, allowing this nutrient to exclusively support the metabolism of T cells. We show that enabling engineered T cells to overcome the glucose restriction of the tumor microenvironment allows for an increased capacity to clear tumors. Finally, we show that this capability can be coupled with human chimeric antigen receptor T cells (CAR-T) to more effectively kill tumor cells in low glucose environments.

## Results

### Heterologous and Functional Expression of Cellobiose Catabolism Proteins in Primary T cells

To impart the ability to catabolize cellobiose as a fuel source for glycolysis, only two proteins are needed: 1) a transporter of cellobiose from the extracellular to intracellular space; and 2) a β-glucosidase that can efficiently hydrolyze the β-1,4 bond that joins the glucose molecules in cellobiose. We chose to work with a pair of previously characterized proteins from *Neurospora crassa*, namely, the cellobiose transporter CDT-1 and the cellobiose glucosidase GH1-1 (**Fig. 1a**), which were reported to enable cellobiose catabolism through heterologous expression into yeast^15^.

**Fig. 1.**
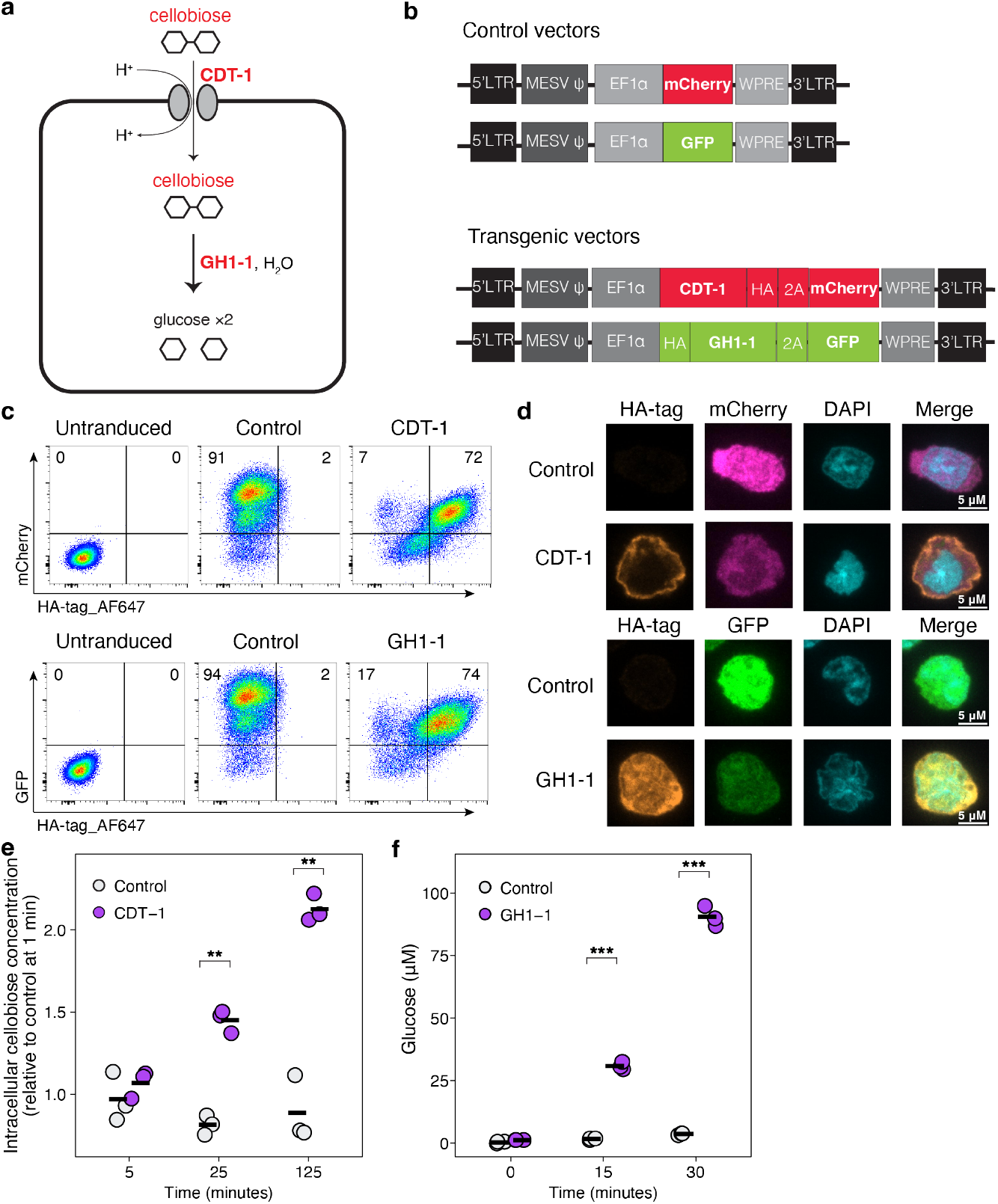
Heterologous expression of two proteins from fungi allow for cellobiose import and hydrolysis in T cells. **a)** Schematic of cellobiose transport through the proton symporter CDT-1, followed by hydrolysis by the cellobiose glucosidase GH1-1. Once glucose is liberated within the cell it may flow through the standard glycolytic pathway. **b)** Schematic of mouse stem viral vectors. Control vectors express fluorescent protein alone and transgenic vectors couple transgenic expression to fluorescent protein expression with a 2A peptide motif. Expression is driven by a constitutive EF1a promoter and enhanced with a woodchuck hepatitis virus post-transcriptional regulatory element (WPRE). **c)** Transgene and fluorescent protein expression analysis in transduced, primary, mouse T cells. **d)** Confocal microscopy images showing localization of CDT-1 (top) and GH1-1 (bottom), fluorescent proteins, and nuclei (DAPI) in transduced, primary, mouse T cells. **e)** CDT-1 transport assay. Control and CDT-1 transduced mouse T cells were incubated with 5 mM cellobiose before extracting intracellular metabolites for LC-MS quantification. **f)** GH1-1 hydrolysis assay. Control and GH1-1 transduced mouse T-cell lysates were incubated with 1 mM cellobiose before quantification of glucose. Horizontal lines indicate means. Statistical significance was assessed using an unpaired t-test in E and F (*p < 0.05; **p < 0.01; ***p < 0.001; ns > 0.05). Data shown in **e** and **f** are means of technical replicates (n=3) from one experiment and are representative of at least two independent experiments.

We optimized the codon sequence of each gene for murine expression (**Fig. S1**) and cloned them into mouse stem cell viral vectors (**Fig. 1b**). Each transgene was appended with a hemagglutinin (HA) tag and coupled to a discrete fluorescent protein via a 2A peptide sequence. Staining of transduced, primary CD8+ T cells for the HA tag revealed robust expression of both of the genes, which was demonstrated to be coupled to their fluorescent reporters (**Fig. 1c**). To confirm the subcellular localization of each protein, we immunostained T cells and performed confocal microscopy. CDT-1 localizes to the periphery of the T cell, which is consistent with its function as a transporter. In contrast, the distribution of GH1-1 is more diffuse, indicative of a cytosolic localization and consistent with the localization of other enzymes involved in the glycolytic pathway^16^ (**Fig. 1d**).

We next sought to confirm the functionality of each transgenic protein. To test the capacity of CDT-1 to transport cellobiose, we incubated transduced T cells with cellobiose, extracted intracellular metabolites, and performed liquid chromatography-mass spectrometry (LC-MS). We found that cellobiose accumulated in the intracellular metabolite pool at a higher rate in T cells singly transduced with CDT-1 compared to control cells (**Fig. 1e**).

To evaluate the functionality of GH1-1 as a cellobiose glucosidase, we singly transduced T cells with GH1-1, generated cellular lysates, incubated with cellobiose, and quantified the resultant glucose yield. We found that GH1-1 expression significantly increased the glucose concentration over time compared to controls (**Fig. 1f**).

Expression of the transgenes in T cells alone did not significantly alter their transcriptional programs (**Fig. S2**). All the metabolic genes examined showed similar transcription between CG-T cells and control-transduced T cells. No Reactome pathways were enriched in CG-T cells as compared with control transduced T cells (neither treated with cellobiose nor low glucose conditions). This result simply shows that transcriptional programs of T cells effectors bearing the CG transgenes are unperturbed.

Taken together, we have developed a system wherein primary T cells co-transduced with the genes CDT-1 and GH1-1 (referred to as CG-T cells) are endowed with the necessary machinery to utilize extracellular cellobiose to provide a source of intracellular glucose.

### Transgenic T cells Robustly Catabolize Cellobiose to Fuel Glycolysis

To test if CG-T cells are capable of catabolizing cellobiose into the appropriate metabolic pathways, we incubated control and CG-T cells with isotopically labelled ^13^C-cellobiose (labeled on all 12 carbons) in the presence of a low concentration of unlabeled glucose (**Fig. 2a**). We observed extensive carbon labeling in key metabolites of the glycolytic, pentose phosphate, and tricarboxylic (TCA) pathways in CG-T cells (**Fig. 2b**), with cellobiose contributing nearly 100% of the carbons present in glycolytic metabolites such as fructose 1,6-bisphosphate, 3-phosphoglyceric acid, and phosphoenolpyruvate. In stark contrast, control cells showed negligible labeling of intracellular metabolites, an expected result that confirms our conventional understanding that mammalian cells lack the capacity to metabolize cellobiose.

**Fig. 2.**
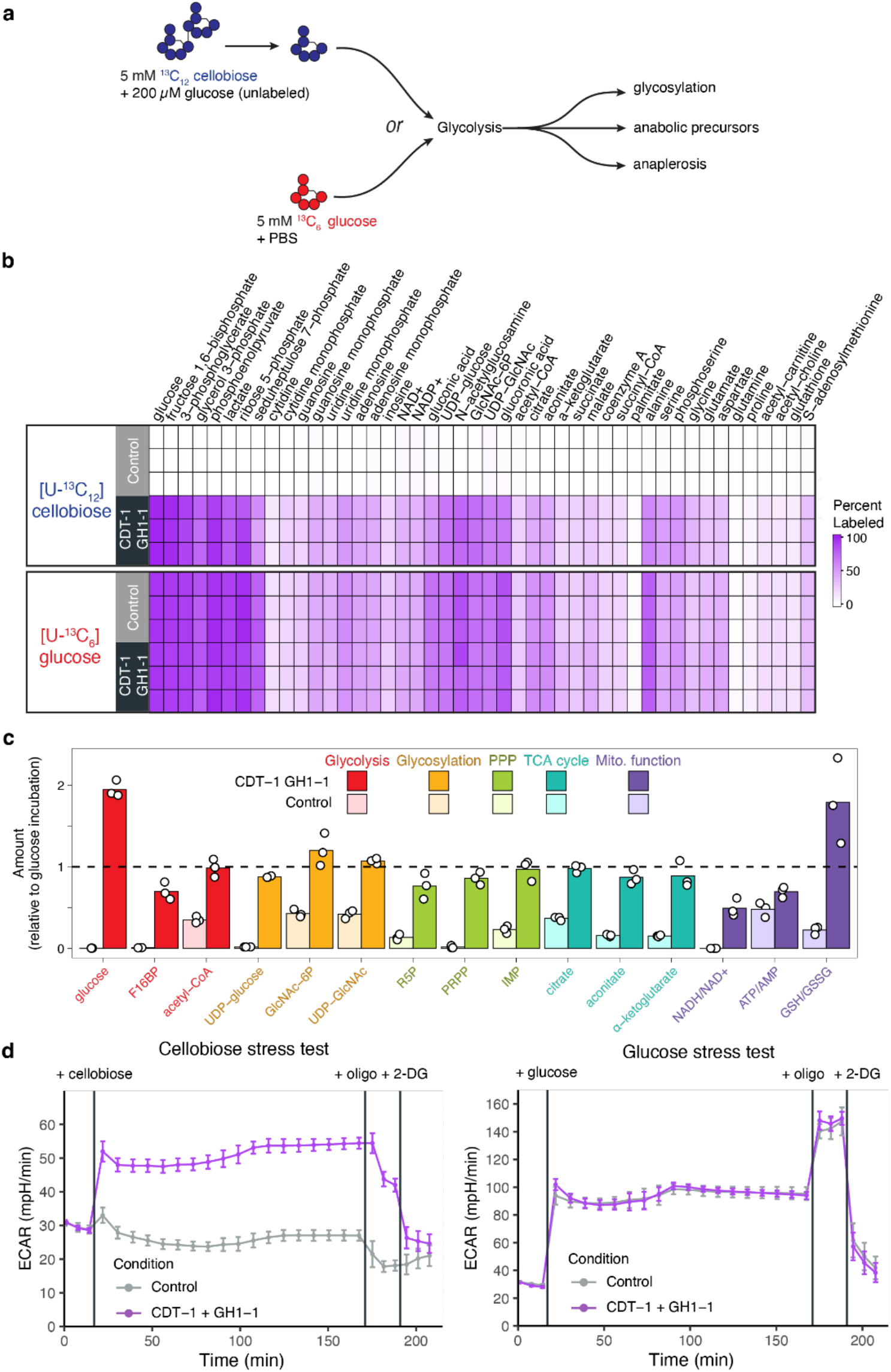
Transgenic T cells metabolize cellobiose and replenish metabolic pathways during glucose withdrawal. **a**) Schematic of isotopically labeled sugar experiment. Transduced T cells were incubated with either 5 mM universally ^13^C-labeled cellobiose and 200 µM unlabeled glucose or 5 mM universally ^13^C-labeled glucose for 16 h before intracellular metabolites were extracted for analysis by LC-MS. **b**) Heat map displaying the percent labeling of intracellular metabolites from transduced T cells incubated with isotopically labeled cellobiose or isotopically labeled glucose. **c**) Abundance of intracellular metabolites during cellobiose incubation relative to their abundance during glucose incubation. PPP, pentose phosphate pathway. **d**) Cellobiose and glucose stress tests using a XFe96 Seahorse Flux Analyzer. Transduced T cells had basal measurements taken before injection of cellobiose or glucose from the first port. Oligomycin A was injected from the second port and 2-deoxyglucose (2-DG) injected from the third port. ECAR, extracellular acidification rate. Error bars represent standard deviation of the mean, n=12. Data shown in b and c are from technical replicates (n=3) of one experiment and are representative of at least two independent experiments.

To determine whether cellobiose carbons are fluxing through the standard glycolytic pathway, we incubated the transduced T cells with isotopically labeled ^13^C-glucose (labeled on all six carbons) and compared the pattern of carbon labeling of glucose metabolites to labeling of cellobiose metabolites. This comparison in CG-T cells revealed a striking similarity in the distribution profile of catabolized cellobiose carbons to glucose carbons (**Fig. 2b**), demonstrating that cellobiose carbons are flowing through the standard metabolic pathways. When comparing glucose catabolism in CG-versus control T cells, the distribution profiles of glucose carbons again look nearly identical, demonstrating that transgenic expression of CDT-1 and GH1-1 did not result in any changes in the fate of catabolized glucose carbons. These results confirm that our approach allows cellobiose catabolism to replace the role of glucose in feeding metabolic pathways.

Glucose deprivation in T cells leads to profound metabolic perturbations, with reductions in the amounts of metabolites from the glycolytic, glycosylation, pentose phosphate, and tricarboxylic acid (TCA) pathways, and the accumulation of low-energy and oxidative intermediates including NAD+, AMP, and oxidized glutathione (GSSG) (**Fig. S3**). To assess whether CG-T cells can use cellobiose to normalize metabolite abundances during glucose withdrawal, we compared the abundance of metabolites during incubation with cellobiose to the abundances during incubation with glucose. CG-T cells incubated with cellobiose showed rescue of core glycolytic metabolites (glucose, F16BP, and acetyl-CoA), intermediates derived from glycolysis that contribute to protein glycosylation (UDP-glucose, GlcNAc-6P, and UDP-GlcNAc), and precursors of purine and pyrimidine synthesis (ribose-5-phosphate, phosphoribosyl pyrophosphate, and inosine monophosphate) (**Fig. 2c**). Furthermore, incubation with cellobiose normalized levels of metabolites central to mitochondrial function, with increased amounts of TCA substrates (acetyl-CoA, citrate, aconitate, and α-ketoglutarate) and the accumulation of a reduced chemical environment (NADH, ATP, and GSH) (**Fig. 2c**). These results demonstrate that in CG-T cells, cellobiose is catabolized into glucose, which then replenishes several of the major metabolic pathways that are depleted during glucose deprivation.

To compare the glycolytic output of CG-T cells using cellobiose versus glucose as fuel sources, we conducted sugar stress tests using the Seahorse platform, wherein the starting condition entailed complete glucose deprivation and then the first injection was either glucose or cellobiose. CG-T cells exposed to cellobiose led to robust engagement of glycolysis, demonstrated by a ∼2-fold increase in extracellular acidification rate (ECAR). Control T cells did not change ECAR when exposed to cellobiose. As expected, glucose exposure equivalently increased ECAR in both control T cells and CG-T cells (**Fig. 2d**), again demonstrating that the transgenes do not alter the capacity to engage glycolysis. These results confirm the ability to use cellobiose as a fuel source. The discrepancy between ECAR increase seen when CG-T cells were exposed to glucose versus cellobiose may be impacted by the mechanism of CDT-1 transport, namely that a proton is transported into the cell along with cellobiose (**Fig. 1a**). As catabolism of cellobiose generates protons that are excreted into the media, CDT-1 transport of cellobiose removes protons from the media, ultimately diminishing the net readout of extracellular acidification. Together, these findings demonstrate that the profound metabolic disturbances experienced by T cells upon glucose withdrawal can be ameliorated by cellobiose through the expression of CDT-1 and GH1-1.

### Effector Function of Transgenic T cells is Fueled by Cellobiose During Glucose Withdrawal

Activated T cells are critically dependent on glucose for optimal anti-tumor activity, as reductions in glucose negatively impact survival, proliferation, and cytokine production. To validate whether cellobiose could rescue these functions in transgenic T cells during glucose deprivation, we next performed a series of in vitro tests.

To assess whether cellobiose could enhance T-cell survival during glucose withdrawal, we incubated T cells ± glucose ± cellobiose for 48 h. Under light microscopy, control T cells incubated with glucose appeared large, polarized, crawling, and healthy. In contrast, cells exposed to low glucose adopted the distinct morphological characteristics of dead and dying cells – that is they appeared rounded and blebby (**Fig. 3a** and **b**). CG-T cells incubated with and without glucose appeared the same as their control counterparts, and the inclusion of cellobiose during glucose withdrawal rescued vitality and cellular morphology akin to cells incubated with glucose. The presence of cellobiose had no discernible effect on morphology in control cells. Thus, cellobiose extends survival and morphology in CG-T cells during glucose withdrawal. We further assessed the same cell populations and their morphological characteristics by flow cytometry. When incubated with glucose, a high fraction of the control cells was large and had low granularity (FSC^hi^, SSC^lo^) (**Fig. 3c**). The absence of glucose shifted the cell population towards a FSC^lo^, SSC^hi^ phenotype, commonly interpreted as dead and dying compared to their large, low granularity counterparts. Again, the presence of cellobiose made little difference for control cells. In contrast, CG-T cells incubated with cellobiose during glucose withdrawal shared the morphology of cells incubated with glucose, demonstrating that transgenic CG-T cells sustained their viability when using cellobiose as a metabolic substrate.

**Fig. 3.**
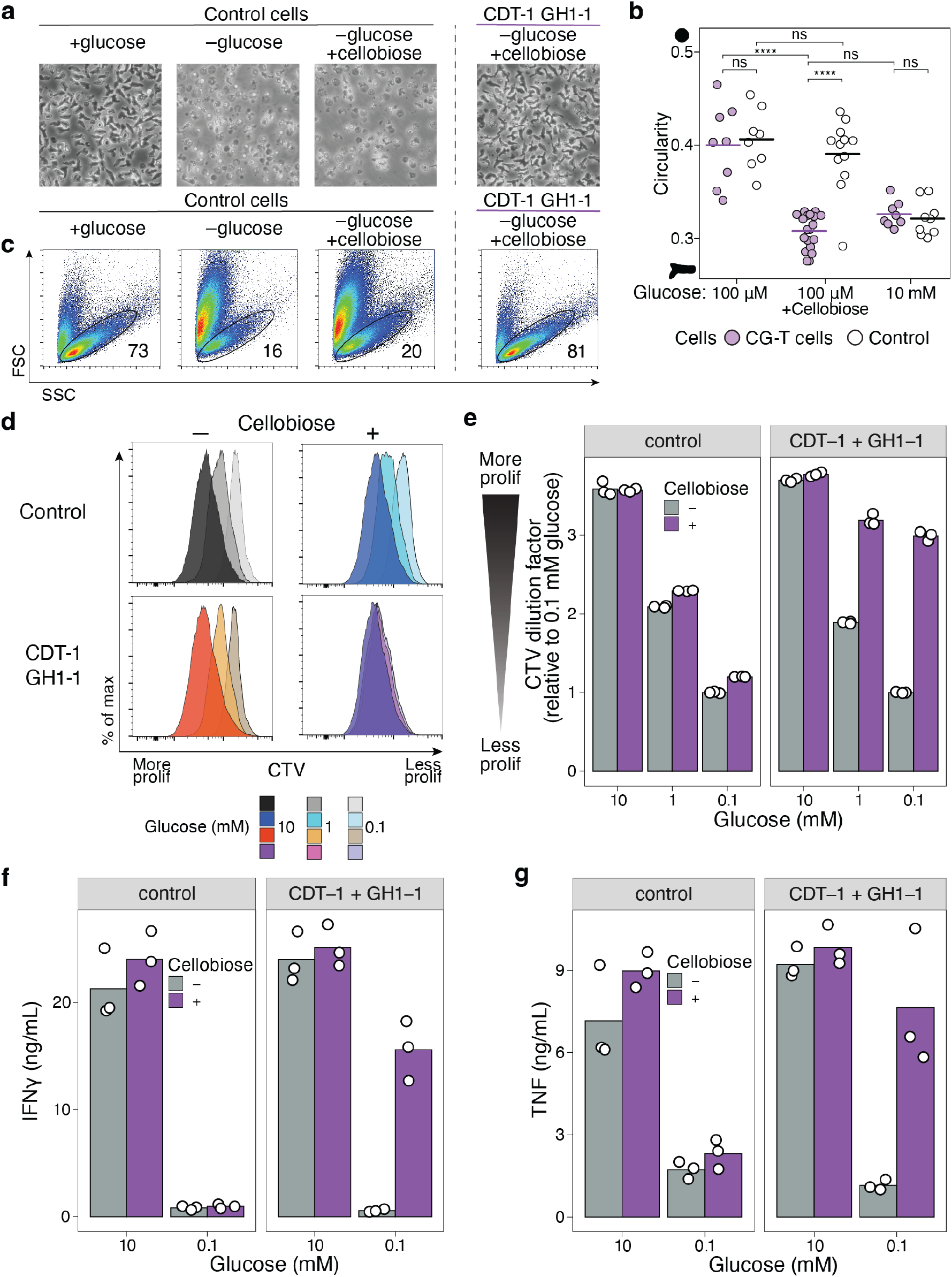
Cellobiose rescues effector functions in transgenic T cells. **a**) Light microscopy of transduced T cells after 48 hours in the indicated metabolic conditions. **b)** Quantification of rounded/circular shape (unhealthy) versus polarized (healthy) in CG-T and control-transduced cells in the indicated metabolic conditions. Horizontal lines indicate means. Each dot indicates the average for many individual T cells within a distinct microscope frame. Statistics shown are t-test. corrected for multiple testing. **c**) Flow cytometry was used to assess size (FSC) and granularity (SSC) of the same cell populations as imaged in (a). **d**) Transduced T cells were stained with CTV and immediately incubated in the indicated metabolic conditions for 48 hours, before measuring the extent of CTV signal dilution. Note that the using CTV to stain and analyze activated, cycling T cells does not produce discrete, halved signal peaks from each cell division (as is typically seen when this reagent is used in the context of naïve T cell activation). **e**) Quantification of the CTV signal dilution factor from (**c**). The CTV signal dilution factor was calculated using the geometric mean of the CTV signal from each population. **f** and **g**) Transduced T cells were incubated in the indicated metabolic conditions for 18 h before stimulation with PMA/ionomycin for an additional 4 h. The supernatants were then collected for measurement of cytokines. Data shown in **e-g** are from technical replicates (n=3) of one experiment and are representative of at least two independent experiments. (*p < 0.05; **p < 0.01; ***p < 0.001; NS > 0.05).

To determine whether cellobiose could not only extend survival but also support the more resource and energy intensive process of cell division, transduced T cells were labeled with the cell proliferation dye CellTrace Violet (CTV). After CTV staining, the cells were incubated for 48 h in high to low concentrations of glucose and cellobiose was added or omitted at each concentration. We assessed proliferation by the dilution of CTV as measured by flow cytometry. As expected, reducing glucose concentration resulted in less dilution of the CTV signal (i.e., less cell proliferation) in control and CG-T cells, but the inclusion of cellobiose allowed only CG-T cells to proliferate robustly, even at the lowest glucose condition tested that otherwise eliminates proliferation (**Fig. 3d**). The proliferation of CG-T cells with cellobiose was comparable to their division seen at the highest concentration of glucose (**Fig. 3e**). Thus, cellobiose robustly supports T cell proliferation in low glucose contexts.

To measure the extent to which cellobiose can fuel cytokine production during glucose withdrawal, CG-T cells were incubated in high or low glucose ± cellobiose overnight and then restimulated with PMA/ionomycin. Glucose withdrawal profoundly reduced the production of IFN-γ and TNF, an effect seen in the percent of cytokine positive cells and the concentration of the cytokines in the supernatant (**Fig. 3f** and **g**; **Fig. S4**). During glucose withdrawal, CG-T cells that were incubated with cellobiose restored IFN-γ and TNF production. Thus, cellobiose fuels cytokine production in low glucose contexts.

Taken together, we show that two major effector functions of T cells—proliferation and cytokine production—are rescued by cellobiose when glucose is critically low in T cells transduced with CDT-1 and GH1-1.

### In vivo proof of concept

Many diverse metabolic perturbations develop in cancer cells, affecting the extent to which glucose is depleted from their TME. As a proof of concept for our approach, we sought a suitable model system of cancer in mice that conferred a low glucose microenvironment where cellobiose-metabolizing T cells could potentially thrive in the presence of cellobiose. We subcutaneously implanted into separate mice three different syngeneic cancer cell lines (EL4-OVA (lymphoma), B16-ND4 knockout (melanoma), and MC38-OVA (adenocarcinoma)). We compared the concentration of glucose in healthy tissues (spleen, peritoneum, and skin) to the concentration in these solid tumors by LC-MS. We found that EL4-OVA and B16-ND4 KO tumors consistently had significantly reduced concentrations of glucose compared to the healthy, adjacent tissues, and that EL4-OVA tumors were more depleted of glucose than all the healthy reference tissues (**Fig. S5a**). All three types of tumors had samples with lower glucose concentrations than the healthy tissue, while MC38-OVA tumors were most heterogenous, with some tumors harboring low glucose levels and other tumors in the range of healthy tissue. Based on these results, we proceeded with the EL4-OVA model.

To determine the extent, if any, that the tumor cells themselves could benefit from the presence of cellobiose as a metabolic substrate, we incubated EL4-OVA cells in media lacking glucose but containing cellobiose and assessed the capacity to survive and divide. EL4-OVA cells were incapable of using cellobiose in vitro to facilitate proliferation (**Fig. S5b**). Furthermore, live/dead staining showed no survival advantage for the tumor cells when cellobiose was provided in the absence of glucose (**Fig. S5c**). These results confirm that cellobiose would exclusively be available to CG-T cells as a fuel source. To analyze any impact of cellobiose on the intestinal microbiome, these tumor-bearing mice fed cellobiose or PBS were

For cellobiose to be available as a fuel source for adoptively transferred T cells, it must be present in the blood in high concentrations. Providing cellobiose in food or water is not a viable means of increasing blood concentration of cellobiose, due to the low rates of para- and transcellular transport of cellobiose across the intestinal epithelium, in addition to the capacity for gut microbiota to metabolize cellobiose. Instead, we opted to deliver cellobiose through repeated intraperitoneal injections. To determine the appropriate dosing schedule of cellobiose, we monitored the concentration of cellobiose in the blood after i.p. injection by LC-MS. We found that after i.p. injection, the concentration of cellobiose went up sharply into the millimolar range and returned back down to baseline within 8 h (**Fig. S5e**). Based on these results we decided to administer multiple cellobiose injections each day after the adoptive transfer of CG-T cells.

As T cells consume cellobiose and generate intracellular glucose, we wanted to be sure that they don’t export any of that glucose, which could then be consumed by tumor cells. We tested this by incubating CG-T cells with cellobiose and without glucose and then measuring the amount of glucose in the media 24 hours later. We observed that 5 mM cellobiose medium alone (with no cells) was reproducibly contaminated with a small but measurable level of glucose (approx. ∼20 µM), which was derived from impurities in the cellobiose reagent. CG-T cells exported no measurable glucose into the medium, comparable to control T cells (**Fig. S5e**). Both cell types showed consumption of the trace amount of glucose contamination. CG-T cells consumed the cellobiose and produced lactate into the extracellular media, while control T cells could not, as expected (**Fig. S5f**). These results confirm that CG-T cells cannot indirectly provide glucose to tumor cells in the tumor microenvironment from the cellobiose they consume.

Finally, we tested the ability of CG-T cells fed with cellobiose to improve tumor clearance in a proof-of-concept experiment (**Fig. 4a**). To provide for antigen-specific targeting of T cells to the tumor, we employed OT-I TCR transgenic T cells that recognize the ovalbumin antigen expressed by EL4-OVA tumors. We implanted EL4-OVA tumors subcutaneously on day 0. We activated OT-I T cells and transduced the CDT-1 and GH1-1 transgenes as above, sorting cells for purity before use. T cells were injected i.v. on mice on day 10. Starting on the day of T cell adoptive transfer, we delivered either cellobiose or phosphate buffered saline (PBS, control) by twice daily i.p. injections in a blinded fashion. We measured the size of the tumors by calipers and assessed for survival as determined by a tumor-size cutoff (see Methods) (**Fig. 4b**). Survival was significantly improved in the mice receiving cellobiose injections (Cox regression, p = 0.029) (**Fig. 4c**). The probability of survival at 30 days was 0.67 for mice treated with cellobiose and 0.25 for PBS. After the transfer of T cells, the median survival was 31 days for mice treated with cellobiose and 16 days for PBS. The hazard ratio for survival was 2.95 (95% CI 1.1 to 7.8). At 4 weeks, progression-free survival (PFS) was five of 12 (42%) for mice receiving cellobiose and 8% for control-treated mice. Two of 12 (17%) mice receiving cellobiose showed a complete response whereas none of the PBS-treated mice survived long-term. These results show that tumor growth is more heavily suppressed and survival is prolonged in mice when treated with antigen-specific T cells capable of metabolizing cellobiose.

**Fig. 4.**
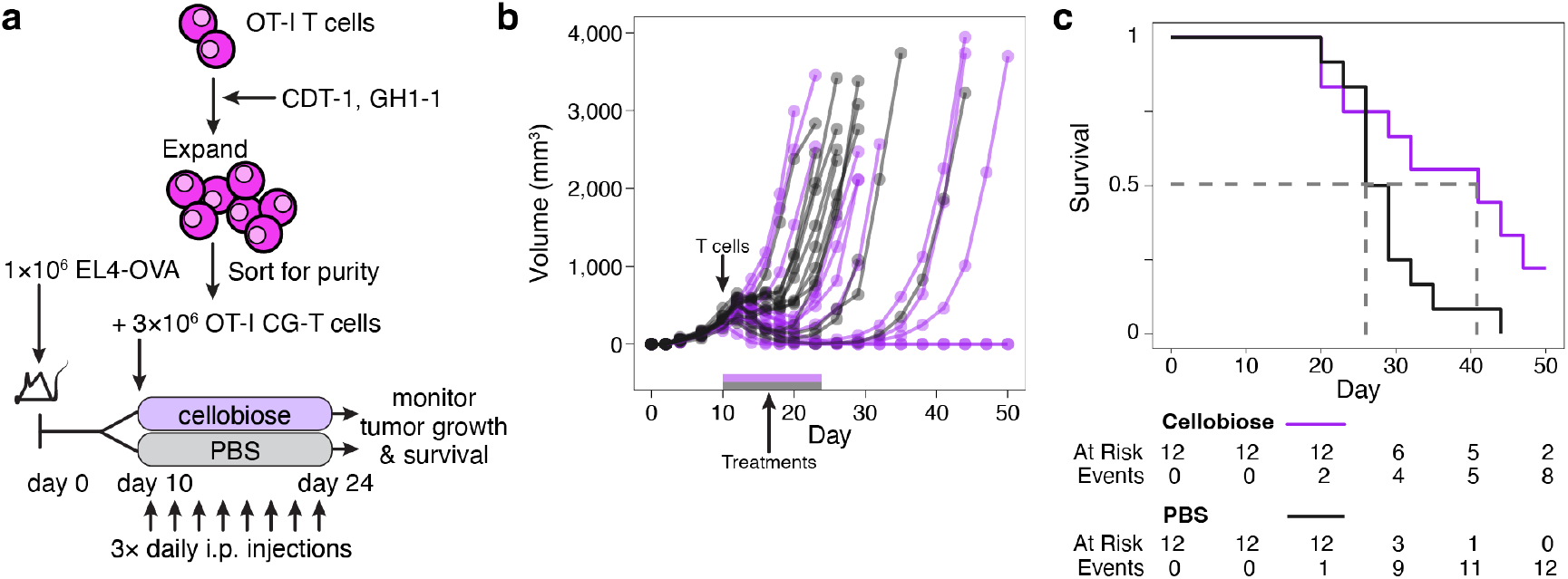
T cells that can metabolize cellobiose are a more effective treatment for cancer. **a**) Schematic for the cancer experiments. **b**) Tumor volumes for mice receiving T cells on day 10 and twice daily i.p. injections of cellobiose (n=12) or PBS (n=12). Survival was defined by tumors reaching a size cutoff (see Methods). **c**) Survival curves show median survival after transfer with T cells of 31 days for mice treated with cellobiose and 16 days for mice treated with PBS. Two of twelve mice treated with cellobiose showed a complete response. This experiment is representative of two.

We next sought to assess the transcriptional programs of CG TILs in the TME (experimental schematic in **Fig. 5a**). We implanted EL4-OVA tumors subcutaneously in Rag1 KO mice on day 0. We transduced OT-I T cells with either the CDT-1 and GH1-1 transgenes (also GFP and mCherry+) or transduced with control vectors (GFP+), expanded and sorted for purity. The T cells were mixed 1:1 and the ratio confirmed by flow cytometry (51% CG-T: 49% control-transduced T cells). By mixing T cells in the same TME, we could identify cell-intrinsic differences conferred by cellobiose metabolism. The tumors were removed five days later, tumors disaggregated, cells pooled, CD8+ T cells enriched by positive-selection magnetic beads, and analyzed by 10x single-cell RNAseq. We found the tumor-infiltrating CD8+ T cells in a variety of states including bearing transcriptional markers of naïve-like, activated, and exhausted states (**Fig. 5b**). Among the single cells analyzed, we identified the CG T cells by their transcription of the transgenes or the mCherry fluorescent protein and found they occupied all the CD8+ states but mostly “naïve-like” and “activated” with only a few exhausted (**Fig. 5b**). Differentially-expressed transcripts in CG-T cells (as compared to control T cells) were analyzed for enrichment of biological processes and found to express transcriptional programs of proliferation, activation, and receptor signaling (**Fig. 5c**). To understand the proliferative aspect better, transcriptional signatures of cell cycles phases G1, G2m, and S were analyzed using Seurat. We found that CG cells expressed more S phase transcripts and fewer G1 transcripts. These results confirms that cellobiose endows CG-TIL with cell-intrinsic enhancement of effector programs, especially proliferation.

**Fig. 5.**
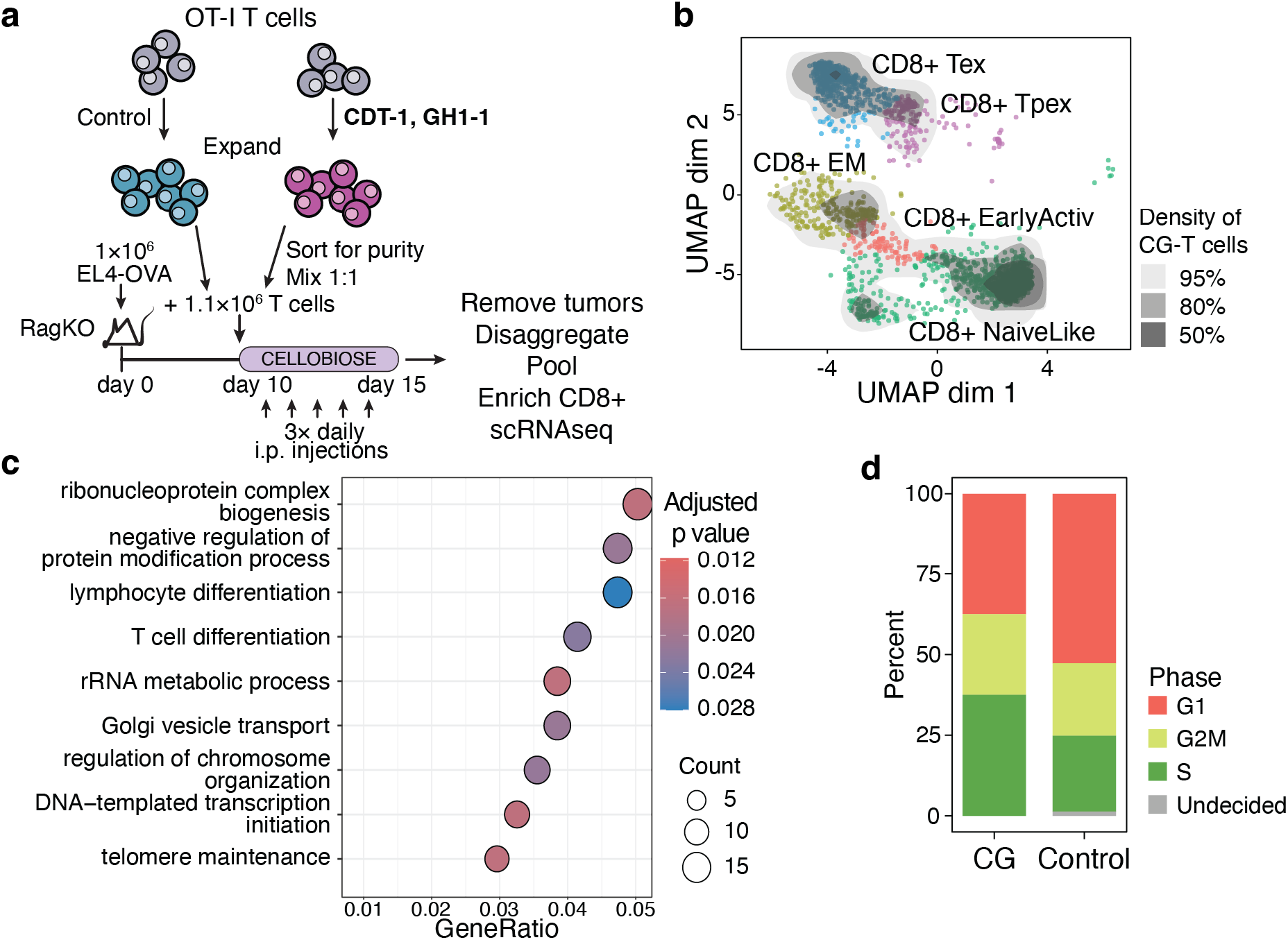
Transcriptional program of tumor infiltrating CG-T cells shows improved effector functions. **a)** schematic of the experiment with co-transfer of CG transgenic and control T cells. **b)** Cell types identified by single-cell RNAseq. The CG T cells are superimposed by density clouds atop the total distribution of identified cell types. 50% of the CG-T cells are found in the darkest regions. T cell types identified as CG by their transcription of the CG transgenes or mCherry fluorescent protein. **c)** GO terms enriched in the CG-T cells compared to non-CG cells. **d)** Proportion of CG and control T cells Cell cycle, calculated by Seurat.

### Human CAR-T cells

Chimeric antigen receptor (CAR)-T cells have arisen as an important immunotherapeutic strategy for a variety of hematogenous tumors. The application of CAR-T cells to solid tumors, on the other hand, has foundered, at least partially due to the reduced glucose environment in the TME^17^. To extend our approach to offer metabolic support to human T cells, here we performed experiments using the model system of human T cells transduced with a standard anti-CD19-CAR construct that targets human CD19, expressed abundantly on Raji tumor cells. T cells were also co-transduced with GH1-1 and CDT-1 viruses, expanded, and flow sorted for the presence of all three proteins (**Fig. 6a**). These experiments were performed along with control T cells that were transduced with only the CD19-CAR.

**Fig. 6.**
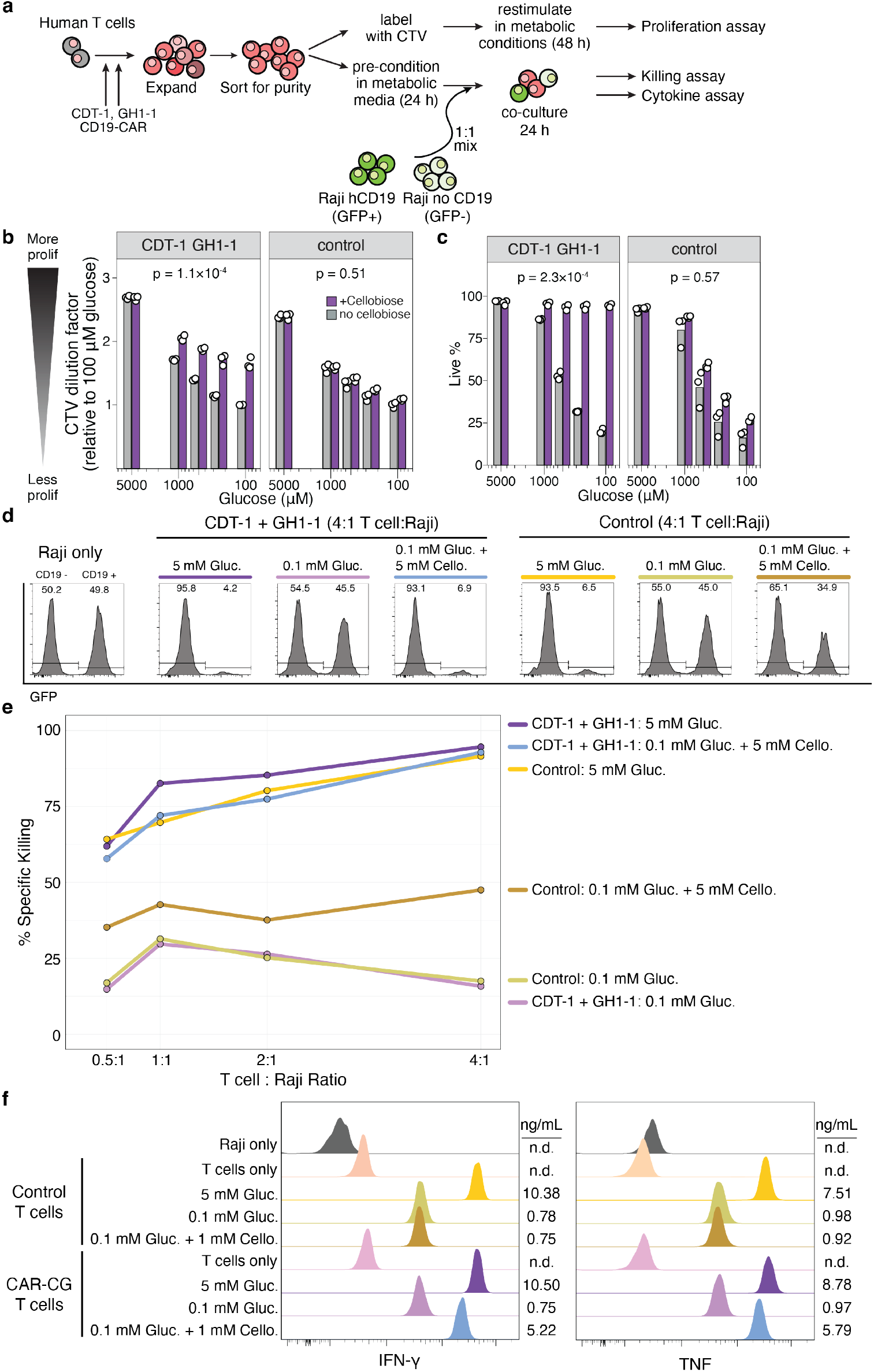
Cellobiose rescues human CAR-T cell function in low glucose conditions. **a**) Schematic of the experiment where human T cells are transduced with CD19 CAR construct plus CDT-1 and GH1-1, or controls expressing the CAR only, and cultured in metabolic conditions for proliferation, killing, or cytokine assays. In killing and cytokine assays, the cells were co-cultured with a 1:1 mixture of CD19+GFP+ and CD19-GFP-Raji B cells at various ratios. **b** and **c)** Proliferation and live staining of CAR-CG-T cells versus CAR-T cells (facets) in low glucose conditions, with (purple) or without cellobiose (gray). Cell trace violet dilutions were normalized to the average of each cell type’s proliferation in low glucose conditions (100 µM) without cellobiose. Two-way (glucose levels and cellobiose) ANOVA p-value shown. Each dot is an independent transduction. **d)** Flow cytometric killing assay showing CAR-T cells (either CG-CAR or CAR alone) co-cultured with Raji cells bearing CD19 (GFP+) and lacking CD19 (GFP–) to assess specific killing. Experiments were performed at a 4:1 T cell:Raji ratio. **e)** Killing is potentiated by cellobiose under low glucose conditions in CDT-1 + GH1-1 CAR T cells. Specific killing of Raji-CD19 cells is shown at various ratios of CAR-T cells (either CG-CAR or CAR alone) to target cells. **f)** Pre-conditioned CAR-T cells were incubated in metabolic conditions with CD19+ Raji B cells at a 1:2 T cell:Raji ratio. IFN-γ and TNF were measured in the supernatants after 24 h.

To ascertain if cellobiose could potentiate proliferation of human CAR-T cells in low glucose environments, we expanded the CAR-T cells, rested, labeled with CTV, restimulated the cells for 48 h, and measured relative proliferation by dilution of CTV. CAR-CG-T cells showed enhanced proliferation in low glucose conditions when cellobiose was provided (**Fig. 6b**). Control (CAR only) T cells and CAR-CG-T cells showed poor proliferation in low glucose conditions alone. CAR-CG-T cells also showed total rescue of viability in low glucose conditions when cellobiose was provided (**Fig. 6c**). Improvements in proliferation and viability were seen across a wide range of physiologically relevant, low glucose conditions. These results copy those previously shown for mouse T cells. Thus, cellobiose potentiated proliferation of human CAR-CG-T cells in low glucose.

To ascertain if cellobiose could improve tumor killing function in low glucose environments, we co-cultured CAR-CG-T cells or control transduced CAR-only T cells with differentially labeled Raji cells, either expressing CD19 (GFP+) or not expressing CD19 (GFP–). Specific lysis of CD19-bearing Raji cells was assessed by flow cytometry, gating on human CD20 (**Fig. 6d**). Across a range of ratios of effector T cells to target cells, we found complete rescue of specific lysis of Raji-CD19 cells in low glucose conditions when cellobiose was provided to CAR-CG-T cells (**Fig. 6e**). Control transduced CAR-T cells showed no benefit from cellobiose and showed profound reduction in killing at low glucose concentrations. When not provided cellobiose, CAR-CG-T cells similarly showed poor killing in low glucose conditions. We found that 5 mM cellobiose contained enough glucose contamination to partially afford some rescue to control T cells. Thus, cellobiose potentiated human CAR-CG-T cell killing in low glucose.

To ascertain the ability of cellobiose to rescue cytokine function in low glucose conditions, we again cultured CAR-CG-T cells or CAR-only T cells in low glucose conditions (0.1 mM) for 24 h, then mixed with Raji target cells at a ratio of 1:2 in culture media comprising various glucose and cellobiose concentrations. We assessed cytokine production from the supernatants by cytokine capture beads. We found dramatic rescue of production of IFN-γ and TNF from human CAR-CG-T cells when provided cellobiose in the context of low glucose conditions (**Fig. 6f**). Control-transduced T cells revealed poor function in low glucose and no rescue with cellobiose, as expected. Accentuation by cellobiose of cytokine production occurred even at very low glucose concentrations, but showed diminishing accentuation as glucose levels rose to 400 µM or higher (**Fig. S6**). These results copy those previously shown for mouse T cells. Thus, cellobiose potentiated human CAR-CG-T cell cytokine production in low glucose.

To study in vivo responses, we injected Raji-hCD19 cells subcutaneously in NSC mice, allowed them to grow for 10 days, re-assorted the mice so that the average tumor size was the same between two groups, then injected mice repeatedly with either cellobiose or PBS and compared the tumor sizes. We saw no differences in average tumor sizes over 10 days (**Fig. S7a**). Thus, Raji tumors do not directly or indirectly benefit from cellobiose. We injected Raji-hCD19 cells subcutaneously in NSC mice, allowed them to grow, and assessed glucose in the tumor microenvironment by LC-MS. We found glucose levels two orders of magnitude lower than healthy tissues (**Fig. S7b**). Thus, Raji tumors when injected subcutaneously, like many other human solid tumors, engender low glucose levels.

To assess the impact of cellobiose on tumor-infiltration and proliferation by CAR-T cells, we injected Raji-hCD19 cells subcutaneously in NSC mice and re-assorted the mice so that the average tumor size was the same between two groups. We then injected CAR-CG-T cells intravenously into all mice. Half the mice received three-times daily cellobiose injections and half received PBS injections. Note, there are no other T cells in these mice than the ones we injected. We took out tumors and measured the penetration and accumulation of T cells in the tumor by immunohistochemistry (**Fig. S8a**). We measured the proliferation of tumor infiltrating T cells by co-expression of Ki67 in individual cells that stained positively for CD3. Within the tumor, large regions of CAR-T cells juxtaposed against large regions of tumor cells (**Fig. S8b** and **Fig. S8c**, for the cellobiose and PBS treated mice, respectively). We examined a zone of ∼0.5 mm at the interface between these two regions and within the tumor where individual T cells rather than sheets of T cells could be found. In this region, there were many individual tumor-infiltrating T cells (**Fig. S8b** and **Fig. S8c**). To measure proliferation and penetration, we picked many non-overlapping regions each ∼0.6 mm^2^ and examined therein the TILs and their Ki67 staining. We found quadruple the proportion of TILs expressed Ki67 in the mice receiving cellobiose versus PBS (43% versus 9.8%, p = 5×10^-6^) (**Fig. S8d**). The density of TILs was comparable in the tumor where cellobiose was given versus PBS (175 T cells / µm^2^ versus 114 T cells / µm^2^, p = 0.18). This experiment shows that human CAR-CG-T TILs are more proliferative in the presence of cellobiose in the tumor microenvironment.

Finally, we tested if the cellobiose-augmented T cells could impact tumor growth. We injected Raji-luc-hCD19 cells and CAR-CG-T cells in NSC mice as above, and some mice were left untreated to monitor unimpeded tumor growth. Mice were either given cellobiose or PBS as above. Tumor size was monitored by luciferase imaging (**Fig. S8e**). Three mice were excluded that either died during anesthesia for luciferase imaging after the second timepoint or never showed tumor growth. We stopped on day 16 when the untreated mice required euthanasia. We saw a trend towards smaller tumor sizes in cellobiose-treated mice with a 9.5-fold difference in median tumor size (p = 0.12 by Wilcox) (**Fig. S8f**). Four of five (80%) cellobiose-treated mice showed total elimination of tumor. Two of the four (50%) PBS-treated mice also showed complete responses (**Fig. S8g**). This experiment showed human CAR-CG-T cells given cellobiose showed a trend towards improved regression and smaller tumors.

Taken together, these experiments show that the CG genes and cellobiose can enhance the ability of human CAR-T cells to proliferate, produce cytotoxic cytokines, and penetrate tumors in a low glucose environment. These results offer evidence of the therapeutic potential of T cells endowed with the ability to metabolize cellobiose.

## Discussion

Engineered T cells (e.g., CAR-T) already play a major clinical role in fighting lymphomas. In contrast, engineered T cells have struggled to make clinical headway against solid tumors. In the solid TME, TILs (including endogenous and engineered T cells) may be directed to the correct targets but are outcompeted by the tumors for a limited supply of glucose, resulting in a stunted anti-tumor response. The upregulation of glucose transporters such as GLUT1/4 and the kinase HK2 in solid tumors result in the voracious uptake of glucose, sequestering this important nutrient away from the extracellular space^18^. Diminishment of glucose at the tissue scale in tumors can exceed ten-fold compared to healthy, adjacent tissues^19^, as our own results substantiated. The impact on the immune response is profound: GLUT1 expression in human tumors negatively correlates with CD8+ T cell infiltration^20^. Regardless of GLUT1 expression, CD8+ TILs showed in one study of human tumors poor glucose uptake and consequently low levels of activation and proliferation^21^. By engineering the ability to metabolize the natural glucose disaccharide cellobiose into antigen-specific T cells, our approach offers the potential to complement cancer immunotherapies like CAR-T cells in their fight against solid tumors.

There are multiple mechanisms by which provision of glucose enhances T cell responses. Glycolysis fuels the biosynthesis of nucleic acids, lipids, and amino acids. These anabolic processes are critical for cellular maintenance, cytokine production, and proliferation. We showed that carbons derived from cellobiose in our engineered T cells precisely fill those biosynthetic pathways in the absence of other sources of glucose. Deprivation of glucose inhibits expression of IFNγ in effector T cells through regulation at the epigenetic, transcriptional, and translational levels. For example, high concentrations of acetyl-CoA (which can be formed from pyruvate, a product of glycolysis) increase histone acetylation, promoting transcription of proinflammatory genes like *Ifng* and *Tnf* ^22^. Additionally, certain glycolytic metabolites, like glyceraldehyde-3-phosphate, can indirectly modulate T-cell function by sequestering GAPDH away from the 3’UTR of the *Ifng* gene. This diversion releases suppressive post-transcriptional control over translation^23^. These effects are important for anti-tumor immunity, and our results confirmed that production of IFNγ in CAR-T cells is rejuvenated by cellobiose in glucose-restricted conditions.

Human tumor microenvironments vary in their glucose concentrations but consistently show low glucose and high lactate levels ^18,19,24,25^. In our hands, human CAR-T cells showed poor viability when the *in vitro* glucose concentration was 500 µM or less. These CAR-T cells showed poor cytokine production when the *in vitro* glucose concentration was below 200 µM. Killing similarly was compromised only when glucose levels were 200 µM or lower. On the other hand, proliferation defects showed a graded response across all low glucose concentrations. Accentuation by cellobiose of cytokine production occurred even at very low glucose concentrations, but showed diminishing accentuation as glucose levels rose above 400-500 µM or higher. Thus, cellobiose potentiates CAR-T cells across a range of physiologically relevant low-glucose conditions.

Cellobiose is exceedingly non-toxic and is indeed “generally regarded as safe” by the US FDA. It is routinely added to many foods including infant formula, drinks, candies, and icings. Oral and intravenous delivery in humans has been tested and shown to be safe ^26,27^, with over 90% of excretion is in the urine^27^. The intestine of rodents contains some bacterial and fungal species that can potentially metabolize cellobiose^28^. Human lactase cannot^29^. Importantly, cellobiose ingestion in humans does not alter blood glucose levels^30^. In our hands, we saw no significant changes in the gut fungal or bacterial microbiome of mice treated with cellobiose (**Fig. S9**). Taken together, our findings and these reports indicate cellobiose is safe to develop for therapeutic use as envisioned here. The capacity to metabolize cellobiose has potential for diverse applications beyond immunotherapy.

Engineering microbial cells to break down cellobiose has been explored in industrial contexts^14^, and can be used for the selection of stably transduced cell lines^31^. The ability to use cellobiose as a glycolytic fuel coupled with isotopically labeled cellobiose could also be a powerful tool that may enable researchers to tease apart the metabolic dynamics between cells in mixed populations. For example, if tumor cells were enabled to metabolize cellobiose and co-cultured with T cells in the presence of isotopically labeled cellobiose, it might be possible to explore which cellobiose-derived (and by virtue glucose-derived) metabolites the tumor cells were producing that accumulate in and affect T cells (S. Kaech, personal communication). Another advantage of this approach for cancer immunotherapy is a built-in kill switch. Many CAR-T cell responses in lymphomas are overactive in the first week and result in morbidity and mortality due to cytokine-release syndrome. If CAR-T cells fueled by cellobiose were to become overactive, the cellobiose infusions could be reduced or withdrawn to curtail cytokine production. Beyond cancer immunology, we believe the development of this method enabling unique access to a keystone nutrient could be used to answer questions about the regulation of metabolism across many cell types, biological processes, and diseases.

## Acknowledgments

We thank Dr. Erika Pearce for her gift of the EL4-OVA cell line. We thank Dr. Greg Delgoffe for advice and for his gift of the B16-ND4 KO cell line. We thank Dr. Yvonne Chen for the gift of the Raji cell lines and the CD19-CAR construct. We thank Michael Trump and Humza Khan for assistance with pilot experiments, and other members of the Butte lab for support. We thank Johanna ten Hoeve for her assistance with LC-MS through the UCLA Metabolomics Core. We appreciate Xinmin Li and the UCLA Technology Center for Genomics & Bioinformatics for sequencing services. We appreciate the staff of the UCLA Translational Pathology Core Laboratory for histology services. We appreciate the services of LumiGenics LLC for assistance with in vivo metabolite quantification and Zymo Research for assistance with microbiome studies.

## Author contributions

Conceptualization: MLM, MJB

Investigation: MLM, RN, WEZ, MAML, TJT

Visualization: MLM, MJB

Funding acquisition: MLM, MJB

Supervision and project administration: MJB

Writing – original draft: MLM, MJB

Writing – review & editing: everyone

## Data availability

All data are available in the main text or the supplementary materials. Sequencing data are available at https://figshare.com/s/d70355c2b27c06fd78b5 (this link will switch to a permanent DOI upon publication)

## Supplementary Materials

Materials and Methods

Figs. S1 to S9

## References and Notes

1. Bantug, G.R., Galluzzi, L., Kroemer, G., and Hess, C. (2017). The spectrum of T cell metabolism in health and disease. Nature Reviews Immunology. 10.1038/nri.2017.99.

2. Ma, E.H., Verway, M.J., Johnson, R.M., Roy, D.G., Steadman, M., Hayes, S., Williams, K.S., Sheldon, R.D., Samborska, B., Kosinski, P.A., et al. (2019). Metabolic Profiling Using Stable Isotope Tracing Reveals Distinct Patterns of Glucose Utilization by Physiologically Activated CD8+ T Cells. Immunity 51, 856-870.e5. 10.1016/j.immuni.2019.09.003.

3. Palmer, C.S., Ostrowski, M., Balderson, B., Christian, N., and Crowe, S.M. (2015). Glucose metabolism regulates T cell activation, differentiation, and functions. Frontiers in Immunology 6, 1–6. 10.3389/fimmu.2015.00001.

4. Peng, M., Yin, N., Chhangawala, S., Xu, K., Leslie, C.S., and Li, M.O. (2016). Aerobic glycolysis promotes T helper 1 cell differentiation through an epigenetic mechanism. Science 1002967, aaf6284–aaf6284. 10.1126/science.aaf6284.

5. Chang, C.-H., Curtis, J.D., Maggi, L.B., Faubert, B., Villarino, A.V., Sullivan, D.O.’, Ching, S., Huang, - Cheng, Van Der Windt, G.J.W., Blagih, J., et al. (2013). Posttranscriptional Control of T Cell Effector Function by Aerobic Glycolysis. Cell 153, 1239–1251. 10.1016/j.cell.2013.05.016.

6. Rolf, J., Zarrouk, M., Finlay, D.K., Foretz, M., Viollet, B., and Cantrell, D.A. (2013). AMPKα1: A glucose sensor that controls CD8 T-cell memory. European Journal of Immunology 43, 889–896. 10.1002/eji.201243008.

7. Gullino, P.M., Clark, S.H., and Grantham, F.H. (1964). THE INTERSTITIAL FLUID OF SOLID TUMORS. Cancer Res 24, 780–794.

8. Ho, P.C., Bihuniak, J.D., MacIntyre, A.N., Staron, M., Liu, X., Amezquita, R., Tsui, Y.C., Cui, G., Micevic, G., Perales, J.C., et al. (2015). Phosphoenolpyruvate Is a Metabolic Checkpoint of Anti-tumor T Cell Responses. Cell 162, 1217–1228. 10.1016/j.cell.2015.08.012.

9. Chang, C.H., Qiu, J., O’Sullivan, D., Buck, M.D., Noguchi, T., Curtis, J.D., Chen, Q., Gindin, M., Gubin, M.M., Van Der Windt, G.J.W., et al. (2015). Metabolic Competition in the Tumor Microenvironment Is a Driver of Cancer Progression. Cell 162, 1229–1241. 10.1016/j.cell.2015.08.016.

10. Andersen, D.W., Daabees, T.T., Applebaum, A.E., Filer, L.J., and Stegink, L.D. (1983). Utilization of intravenously administered β-cellobiose and maltose by young pigs. Journal of Nutrition 113, 1039–1045. 10.1093/jn/113.5.1039.

11. Nakamura, S., Oku, T., and Ichinose, M. (2004). Bioavailability of cellobiose by tolerance test and breath hydrogen excretion in humans. Nutrition 20, 979–983. 10.1016/j.nut.2004.08.005.

12. Lambertz, C., Garvey, M., Klinger, J., Heesel, D., Klose, H., Fischer, R., and Commandeur, U. (2014). Challenges and advances in the heterologous expression of cellulolytic enzymes: A review. Preprint at BioMed Central Ltd., 10.1186/s13068-014-0135-5 https://doi.org/10.1186/s13068-014-0135-5.

13. Bischof, R.H., Ramoni, J., and Seiboth, B. (2016). Cellulases and beyond: The first 70 years of the enzyme producer Trichoderma reesei. Microbial Cell Factories 15. 10.1186/s12934-016-0507-6.

14. Galazka, J.M., Tian, C., Beeson, W.T., Martinez, B., Glass, N.L., and Cate, J.H.D. (2010). Cellodextrin transport in yeast for improved biofuel production. Science 330, 84–86. 10.1126/science.1192838.

15. Ha, S.J., Galazka, J.M., Joong Oh, E., Kordić, V., Kim, H., Jin, Y.S., and Cate, J.H.D. (2013). Energetic benefits and rapid cellobiose fermentation by Saccharomyces cerevisiae expressing cellobiose phosphorylase and mutant cellodextrin transporters. Metabolic Engineering 15, 134–143. 10.1016/j.ymben.2012.11.005.

16. Jang, S., Nelson, J.C., Bend, E.G., Rodríguez-Laureano, L., Tueros, F.G., Cartagenova, L., Underwood, K., Jorgensen, E.M., and Colón-Ramos, D.A. (2016). Glycolytic enzymes localize to synapses under energy stress to support synaptic function. Neuron 90, 278–291. 10.1016/j.neuron.2016.03.011.

17. Peng, J.-J., Wang, L., Li, Z., Ku, C.-L., and Ho, P.-C. (2023). Metabolic challenges and interventions in CAR T cell therapy. Sci. Immunol. 8, eabq3016. 10.1126/sciimmunol.abq3016.

18. García-Canaveras, J.C., Chen, L., and Rabinowitz, J.D. (2019). The tumor metabolic microenvironment: Lessons from lactate. Cancer Research 79, 3155–3162. 10.1158/0008-5472.CAN-18-3726.

19. Hirayama, A., Kami, K., Sugimoto, M., Sugawara, M., Toki, N., Onozuka, H., Kinoshita, T., Saito, N., Ochiai, A., Tomita, M., et al. (2009). Quantitative metabolome profiling of colon and stomach cancer microenvironment by capillary electrophoresis time-of-flight mass spectrometry. Cancer Research 69, 4918–4925. 10.1158/0008-5472.CAN-08-4806.

20. Singer, K., Kastenberger, M., Gottfried, E., Hammerschmied, C.G., Büttner, M., Aigner, M., Seliger, B., Walter, B., Schlösser, H., Hartmann, A., et al. (2011). Warburg phenotype in renal cell carcinoma: High expression of glucose-transporter 1 (GLUT-1) correlates with low CD8+ T-cell infiltration in the tumor. International Journal of Cancer 128, 2085–2095. 10.1002/ijc.25543.

21. Siska, P.J., Beckermann, K.E., Mason, F.M., Andrejeva, G., Greenplate, A.R., Sendor, A.B., Chiang, Y.-C.J., Corona, A.L., Gemta, L.F., Vincent, B.G., et al. (2017). Mitochondrial dysregulation and glycolytic insufficiency functionally impair CD8 T cells infiltrating human renal cell carcinoma. JCI Insight 2, e93411. 10.1172/jci.insight.93411.

22. Peng, M., Yin, N., Chhangawala, S., Xu, K., Leslie, C.S., and Li, M.O. (2016). Aerobic glycolysis promotes T helper 1 cell differentiation through an epigenetic mechanism. Science 1002967, aaf6284. 10.1126/science.aaf6284.

23. Chang, C.-H., Curtis, J.D., Maggi, L.B., Faubert, B., Villarino, A.V., O’Sullivan, D., Huang, S.C.-C., van der Windt, G.J.W., Blagih, J., Qiu, J., et al. (2013). Posttranscriptional Control of T Cell Effector Function by Aerobic Glycolysis. Cell 153, 1239–1251. 10.1016/J.CELL.2013.05.016.

24. Reznik, E., Luna, A., Aksoy, B.A., Liu, E.M., La, K., Ostrovnaya, I., Creighton, C.J., Hakimi, A.A., and Sander, C. (2018). A Landscape of Metabolic Variation across Tumor Types. Cell Systems 6, 301-313.e3. 10.1016/j.cels.2017.12.014.

25. Goveia, J., Pircher, A., Conradi, L.-C., Kalucka, J., Lagani, V., Dewerchin, M., Eelen, G., DeBerardinis, R.J., Wilson, I.D., and Carmeliet, P. (2016). Meta-analysis of clinical metabolic profiling studies in cancer: challenges and opportunities. EMBO Mol Med 8, 1134–1142. 10.15252/emmm.201606798.

26. EFSA Panel on Nutrition, Novel Foods and Food Allergens (NDA), Turck, D., Bohn, T., Castenmiller, J., De Henauw, S., Hirsch-Ernst, K.I., Maciuk, A., Mangelsdorf, I., McArdle, H.J., Naska, A., et al. (2022). Safety of cellobiose as a novel food pursuant to regulation (EU) 2015/2283. EFSA J 20, e07596. 10.2903/j.efsa.2022.7596.

27. Cobden, I., Hamilton, I., Rothwell, J., and Axon, A.T.R. (1985). Cellobiose/mannitol test: physiological properties of probe molecules and influence of extraneous factors. Clinica Chimica Acta 148, 53–62. 10.1016/0009-8981(85)90300-6.

28. Morita, T., Ozawa, M., Ito, H., Kimio, S., and Kiriyama, S. (2008). Cellobiose is extensively digested in the small intestine by β-galactosidase in rats. Nutrition 24, 1199–1204. 10.1016/j.nut.2008.06.029.

29. Strobel, S., Brydon, W.G., and Ferguson, A. (1984). Cellobiose/mannitol sugar permeability test complements biopsy histopathology in clinical investigation of the jejunum. Gut 25, 1241–1246. 10.1136/gut.25.11.1241.

30. Nakamura, S., Oku, T., and Ichinose, M. (2004). Bioavailability of cellobiose by tolerance test and breath hydrogen excretion in humans. Nutrition 20, 979–983. 10.1016/j.nut.2004.08.005.

31. Teixeira, A.P., Stücheli, P., Ausländer, S., Ausländer, D., Schönenberger, P., Hürlemann, S., and Fussenegger, M. (2022). CelloSelect – A synthetic cellobiose metabolic pathway for selection of stable transgenic CHO cell lines. Metabolic Engineering 70, 23–30. 10.1016/j.ymben.2022.01.001.

