## Supplemental for "Enhancing tumor-infiltrating T cells with an exclusive fuel source"

**Supplementary Materials**

Miller ML, …, Butte MJ. **Enhancing tumor-infiltrating lymphocytes with an exclusive and natural fuel source**

### Materials and Methods

#### *Mouse lines*

Mouse strains were obtained from The Jackson Laboratory: C57BL/6J (JAX 000664) and MHC class I-restricted, $OVA_{257\text{-}264}$-specific TCR transgenic (OT-I, JAX 003831). OT-I male mice were bred with C57BL/6J females and females of the F1 generation were used in all studies where OT-I mice were specified. NCG (NOD-*Prkdc*$^{em26Cd52}$*Il2rg*$^{em26Cd22}$/NjuCrl) mice were obtained from Charles River Labs (Strain 572). All mice were bred and maintained under specific pathogen-free standards by the Division of Laboratory Animal Medicine at UCLA. All mouse experiments were carried out in compliance with UCLA's institutional policy on humane and ethical treatments of animals following protocols approved by the UCLA Animal Research Committee.

#### *Virus production*

To begin viral production, $1 \times 10^6$ Platinum-E cells (Cell Biolabs) were plated into 6-well plates in 2 mL DMEM supplemented with 10% heat-inactivated fetal bovine serum (FBS) (antibiotics were omitted from this media to increase transfection efficiency and reduce toxicity) and incubated in a humidified chamber at 37 °C, 5% CO2. 24 h later, Lipofectamine 3000 (Thermo Fisher Scientific) was combined with 400 femtomoles (~2.5 µg) of plasmid according to manufacturer instructions and added dropwise to the well before returning to the incubation chamber. The next morning, the media was aspirated and replaced with 2.5 mL fresh media and returned to the incubation chamber. 24 h later, the media was harvested, and spun at 500 × *g* for 5 min (to remove cellular contamination). To generate media for co-transduction of two viruses when combined with target cells, supernatants from both control conditions were combined (empty vector-mCherry + empty vector-GFP). Similarly, supernatants from both transgenic conditions (CDT-1 + GH1-1) were combined. Viral supernatants were loaded into Amicon Ultra-Centrifugation Filters with a 100k Dalton molecular weight cut-off (Millipore Sigma) before spinning at 1,000 × *g* for 20 min to concentrate. The ~200 µL of viral media generated from concentration was then brought up to 1 mL using complete T cell media containing 2 µg/mL of soluble anti-CD28 and 50 U/mL of human recombinant interleukin-2 (hrIL-2, BRB Preclinical Repository, National Cancer Institute, NIH). 12-well non-tissue culture treated plates were coated with 20 ng/µL RetroNectin (Takara Bio). The 1 mL of viral media was then overlaid into these wells and centrifuged at 1,000 × *g* for 90 min. To increase transduction efficiency, viral media was left in the wells before overlaying T cell suspensions (below). To further ensure high levels of transduction efficiency, viral supernatants were always used for transduction on the same day that the supernatants were harvested.

#### *Primary T cell culture, transduction, and expansion*

$CD8^+$ T cells were isolated from the spleens and inguinal lymph nodes of female mice between the ages of 8-12 weeks old using an EasySep immunomagnetic negative selection enrichment kit (Stem Cell Technologies). Complete T cell media for our experiments comprised RPMI-1640 supplemented with 10% heat-inactivated FBS, 100 U/mL penicillin/streptomycin, 1 mM sodium pyruvate, 10 mM 4-(2-hydroxyethyl)-1-piperazineethanesulfonic acid (HEPES) buffer, and 0.1% β-mercaptoethanol. Tissue culture-treated 12-well plates were coated with 10 µg/mL anti-CD3 (clone 2C11; BioXCell) overnight at 4 °C then washed. T cells were activated by resuspending at a density of $2 \times 10^6$ cells/mL in complete T cell media additionally containing 2 µg/mL of soluble anti-CD28 (clone 37.51; BioXCell) and 50 U/mL of hrIL-2; 1 mL was plated onto each well, and then incubated in a humidified chamber at 37 °C, 5% $CO_2$. After 24 h, T cells were harvested, and supernatant was collected and saved at 4 °C. T cells were resuspended in an equal volume of fresh complete T cell media with 2 µg/mL soluble anti-CD28 and 50 U/mL of hrIL-2 and 1 mL of suspension was overlaid onto virus-loaded plates (see Virus production above). T cells were "spinfected" by centrifuging the plates at 1,000 × *g* at 32 °C for 90 min. Then the plates were incubated in the 37 °C incubator for an additional 90 min. The viral media was then carefully removed, leaving behind the adherent T cells, before adding back 1 mL of the previously stored supernatant from overnight activation in addition to an extra 1.5 mL of fresh complete T cell media supplemented with 2 μg/mL of soluble anti-CD28 and 50 U/mL of hrIL-2. Transduced T cells were incubated at 37 °C overnight. The next morning, T cells were harvested and transferred into 15 mL of fresh complete T cell media supplemented with 50 U/mL of hrIL-2 before incubating at 37 °C for another 24 h. The following day, T cells were harvested and resuspended in fluorescence associated cell sorting (FACS) buffer (comprising Dulbecco's PBS supplemented with 2% FBS and 1 mM ethylenediaminetetraacetic acid (EDTA)) and sorted for double positive cells (GFP+ mCherry+) using a Sony SH800S flow cytometric sorter. Post-sorting expansion involved daily passaging of T cells into fresh complete T cell media, expanding the total culture volume 4-fold with each daily passage. All experiments were performed on cells 5-7 days after activation.

***Expression cassette design and construction***

To generate the control viral vectors, a plasmid containing the mouse stem cell virus (MSCV) backbone (Addgene #24828) was modified to introduce new expression cassettes containing the desired promoter, cloning site, fluorescent reporter, and regulatory element (Fig. 1a). First, the viral cargo between the truncated gag and 3' long terminal repeat (LTR) was removed by amplifying the plasmid backbone, using Phusion polymerase (Thermo Fisher Scientific) and the primers 5'-GGCGCCTAGAGAAGGAGTG-3' and 5'- ATGAAAGACCCCACCTGTAG-3'. The PCR product was then combined with separate gBlocks Gene Fragments (Integrated DNA Technologies) containing the new expression cassettes and assembled with NEBuilder HiFi DNA Assembly Master Mix (New England Biolabs).

To generate the transgenic viral vectors, the amino acid sequences for cdt-1 (KEGG ID NCU00801) and gh1-1 (KEGG ID NCU00130) were codon optimized for murine expression using the GenSmart Codon Optimization Tool (GenScript) and constructed as gBlocks Gene Fragments (Integrated DNA Technologies) with 25 base pair overlaps at the 5' and 3' ends that provided homology to the control plasmids after digestion with NotI and BamHI.

***Metabolite extraction***

For cells in suspension: cells were harvested, pelleted at 500 × *g* for 60 sec, and washed with ice-cold PBS before adding 1 mL of a mixture of 40:40:20 methanol:acetonitrile:water to each cell pellet. Each sample was then spiked with 1 nmol of Norvaline (Millipore-Sigma) before vigorously vortexing for 30 sec. Samples were then placed at -20 °C for 1 h to aid extraction/ protein precipitation and then again vortexed for 30 sec at the end of the extraction incubation. Samples were then spun at 16,000 × *g*, 4 °C for 10 min to pellet cellular debris. Equal volumes (~900 μL) of supernatants were then transferred into glass vials before drying in an EZ-2 Elite (GeneVac) for 60 min. To determine the cell pellet protein content for normalization, the cell pellet was lysed with 0.2 M sodium hydroxide, heated to 95 °C for 20 min, and then quantified using a Pierce BCA assay (Thermo).

For tissues: ~25 mg of flash frozen tissue was added to a microcentrifuge tube containing a 5 mm grinding ball (440C Stainless Steel, OPS Diagnostics) and 1 mL of an mixture of 80:20 methanol:water. The tissues were then homogenized on a TissueLyser II (Qiagen), using 30 oscillations/sec in pulses lasting 30 sec until the tissue was homogenously broken down. The pellets were then incubated at -80 °C for 1 h to aid extraction and protein precipitation, and then vortexed for 30 sec. Samples were then spun at 16,000 × *g*, 4 °C for 10 min to pellet cellular debris and a volume corresponding to 5 mg of tissue was transferred into a glass vial before drying in an EZ-2 Elite to evaporate the fluid.

***Metabolomics***

Dried metabolites were resuspended in 50% acetonitrile (ACN) in water, and an aliquot was loaded onto a Luna 3 μm NH2 100 A (150 × 2.0 mm) column (Phenomenex). The chromatographic separation was performed on a Vanquish Flex (Thermo Scientific) with mobile phases A (5 mM NH4AcO pH 9.9) and B (ACN) and a flow rate of 200 μL/min. A linear gradient was run from 15% A to 95% A over 18 min and was followed by 7 min isocratic flow at 95% A and re-equilibration to 15% A. Metabolites were detected with a Thermo Scientific Q Exactive mass spectrometer run with polarity switching (+3.5/−3.5 kV) in full scan mode with an *m/z* range of 70–975. The open-source Maven application (version 8.1.27.11) was used to quantify the targeted metabolites by area under the curve using expected retention time and accurate mass measurements (<5 ppm). The cell culture experiments were normalized to protein content. The tissue experiments were normalized to tissue content (weight). Data analysis was performed using in-house R scripts (https://github.com/graeberlab-ucla/MetabR).

***Microscopy***

Wells of Nunc Lab-Tek II Chambered Coverglass (Thermo Fisher) were coated with 100 µg/mL poly-d-lysine in phosphate-buffered saline (PBS), incubated at 4 °C overnight, aspirated, and allowed to air dry. 50,000 cells in 200 µL complete T cell media were gently overlaid in a well of the chamber and allowed to settle in a humidified, 5% $CO_2$ chamber at 37 °C for 1 h. The cells were then washed with PBS and then incubated at room temperature for 30 min with a 4% paraformaldehyde in PBS solution. After two washes with PBS, the cells were incubated with intracellular staining permeabilization wash buffer (BioLegend) for 5 min at room temperature before blocking for 60 min at room temperature using 10% donkey serum diluted in permeabilization buffer. After blocking, the cells were incubated in permeabilization buffer with a 1:200 dilution of anti-HA-tag (clone 16B12, BioLegend) conjugated to Alexa Fluor 647 for 60 min at room temperature. The cells were washed twice with permeabilization buffer, and a third time with PBS, before overlaying Fluoromount-G Mounting Medium with DAPI (Invitrogen). The cells were imaged at 100× magnification using a Ti Eclipse (Nikon) microscope, modified with a CSU-X1 (Yokogawa) confocal scanner unit and a XR/MEGA-10 (Stanford Photonics) camera. Images were acquired using MicroManager and analyzed in Fiji.

***Light microscopy and polarization***

CG-transduced T cells or control-transduced T cells were cultured for 48 hours in metabolic conditions. Light microscopy images of cells in plates were taken with an Infinity monochrome microscopy cameras attached to an upright Nikon microscope. Images underwent processing in Fiji, first subtracting background, then using standard menu commands to smooth and despeckle the image, threshold into black and white regions using the Huang filter, eroding the resulting objects, filling holes, and then analyzing for cell shape using the AnalyzeParticles command. The resulting circularity was analyzed and visualized in R.

***Glucosidase activity assay***

T cells were lysed using Pierce IP Lysis buffer (Thermo) supplemented with Halt protease inhibitor cocktail (Thermo), incubated on ice for 10 min with periodic vortexing, and centrifuged at 15,000 × *g* at 4 °C for 15 min. The protein concentration of the supernatant was then quantified using a bicinchoninic acid (BCA) assay (Pierce Thermo). 10 µL of lysate (corresponding to 50 µg of protein) was then combined with 10 µL cellobiose (final concentration of 1 mM), and 30 µL of reaction buffer (50 mM phosphate buffer, pH 6.0)[1] before overlaying 50 µL of working reagent from a glucose detection assay (Amplex Red Glucose/Glucose Oxidase Assay Kit, Invitrogen). Glucose production and oxidation was detected using a Biotek Cytation5 plate reader to measure absorbance at ~560 nm over time.

***Cellobiose export***

Transduced T cells (CG or control) at a concentration of $2.5\times10^6$ / mL were cultured overnight (18 h) in 5 mM cellobiose. To calculate absolute amounts of glucose, standards were run including included glucose free media and media spiked with 10 µM, 100 µM, and 1 mM $^{13}$C-glucose. We measured glucose and lactate by LC-MS from the culture media. Lactate values are reported as uncalibrated spectral counts.

***Bulk RNAseq***

Transduced T cells (CG or control) were grown in IL-2 until day 5, then counted. $3\times10^6$ T cells were pelleted, RNA prepared and quantified, and 320 million paired-end reads were sequenced from a Novaseq X. 76% of the reads were aligned to a modified mouse genome that included the addition of the transgenes (CDT-1 and GH1-1) and fluorophore genes (mCherry and GFP). The resulting counts were analyzed using R scripts. Differential gene analysis was performed with DeSEQ2 [2]. To examine for any metabolic or functional pathways associated with transcript differences, the list of genes enriched with FDR

< 0.05 was entered into the Reactome analysis pipeline (https://reactome.org/ [3]). No resulting pathway showed a p-value less than 0.05.

***Proliferation assay***

Transduced T cells were stained with 1 µM CellTrace Violet (Thermo Fisher) reagent in PBS with 0.1% BSA and incubated for 15 min at 37 °C with regular agitation. Staining was neutralized with complete T cell media before pelleting cells and resuspending in basal media for metabolic assays, composed of RPMI 1640 minus glucose supplemented with 10% dialyzed, heat-inactivated FBS (Thermo Fisher), 100 U/mL penicillin/streptomycin, 10 mM HEPES buffer, 0.1% β-mercaptoethanol, and 50 U/mL of hrIL-2. An equal volume of basal media supplemented with 2x glucose or cellobiose concentrations was then overlaid to achieve the desired concentration for glucose or cellobiose. The cells were incubated in a humidified chamber at 37 °C, 5% $CO_2$ for 48 h before harvesting for CTV signal analysis by flow cytometry.

***Cytokine production assay***

Transduced T cells were incubated for 18 h in basal media (composed of RPMI 1640 minus glucose supplemented with 10% dialyzed, heat-inactivated FBS (Thermo Fisher), 100 U/mL penicillin/streptomycin, 10 mM HEPES buffer, 0.1% β-mercaptoethanol, and 50 U/mL of hrIL-2) plus or minus the indicated concentrations of glucose or cellobiose. To measure cytokine in the supernatant, the cells were then stimulated with PMA (40 ng/mL) and ionomycin (1 µM) for 6 h, then supernatants were diluted before quantification of cytokines by Cytokine Bead Array (BD Biosciences), following the manufacturer instructions. To measure cytokine production on a per-cell basis, the cells were stimulated with PMA (40 ng/mL) and ionomycin (1 µM) in the presence of Brefeldin A (5 µg/mL) for 6 h. T cells were then fixed in 2% paraformaldehyde and 5% sucrose solution for 20 min at room temperature, before permeabilization and staining for intracellular cytokines, and measurement on a flow cytometer.

***EL4-OVA tumor model***

For tumor experiments, cohorts of female C57BL/6J mice born on the same day and between 8-12 weeks old were used. The physical health and behavior of mice were monitored daily to ensure that they did not lose more than 20% of starting weight, did not show signs of stress, and could continue to feed and drink uninhibited. Nair was sued on the abdominal area to remove hair. The mice were injected subcutaneously in the abdomen (between the inguinal nipples) with $1\times10^6$ EL4-OVA tumor cells on day 0. The tumors were allowed to grow until day 10 (when tumors were ~7-8 mm in diameter), then the mice were re-asssorted into two groups such that the average size of the tumors was the same in both groups. The mice then underwent i.v. injection of $3\times10^6$ OT-I CG-T cells. Starting on the day of T cell adoptive transfer, the mice were injected three-times daily with 85 mg cellobiose i.p. or PBS in the same volume in a blinded fashion until day 24. Tumor sizes continued to be followed until the endpoint. The endpoint for each mouse was determined based on physical health, tumor ulceration (which did not occur for EL4-OVA), or if the tumor exceeded 22 mm in diameter in any dimension, or the average of the tumor length or width exceeded 17 mm. Tumor volumes were calculated as $4/3 \times \pi \times [(L+W)/2\ ]^3$, where L and W are the measurements of length and width by calipers. Thus, the survival cutoffs we used correspond to large tumors, volumes between 2000-4000 $mm^3$. Statistical comparisons were done using the R package `survival`, using the Cox regression command `coxph` using a volume of 2200 $mm^3$ as a cutoff.

***Co-adoptive transfer single-cell RNAseq***

Rag1 KO mice were injected subcutaneously as above with $1\times10^6$ EL4-OVA tumor cells on day 0. Tumors were allowed to grow until day 10 (when tumors were ~7-8 mm in diameter). OT-I T cells were activated and transduced as above and expanded in IL-2 and sorted on the fluorescent proteins for purity, then counted. Exactly equal ratios of CG-T cells and control transduced T cells were mixed, and the resulting mixture was verified by flow cytometry (51% to 49%). The mice then underwent i.v. injection of $1.1\times10^6$ mixed T cells. Mice also underwent injections of anti-PD1 (25 µg/mouse/day x 4 days). Mice were euthanized 5 days later and tumors removed.

Tumors were disaggregated and pooled. Magnetic enrichment of CD8+ T cells was performed using EasySep Mouse CD8a Positive Selection. Flow sorting was then used to enrich for T cells. Of the

Vα2+ T cells indicating the OT-I transgenic TCR, 51% were mCherry+, indicating the CG-T cells. The remainder were (GFP-only) control-transduced T cells. T cells were submitted to the UCLA Technology Center for Genomics & Bioinformatics for 10x sequencing.

Resulting fastq files were processed using kallisto / bustools [4]. The mouse genome Mus_musculus.GRCm39.111.gtf was modified to include the cDNA sequences for mCherry, GFP, CDT-1 and GH1-1. A custom kallisto index was calculated. The kb count command was used to to pseudoalign the reads and bustools used to quantify the data. The resulting matrix of counts for barcodes (cells) versus genes was read into a custom R script that relied heavily on the Seurat package version 5[5]. Reads were normalized, scaled and dimensionally reduced for visualization, first with PCA using 2,000 variable genes. The first 20 components were used as input for UMAP visualization in two dimensions. CG-T cells were identified by expression of any combination of GH1-1, CDT-1, or mCherry transcripts. Cell cycle scores were calculated using the CellCycleScoring function in Seurat with default parameters and human gene names converted to mouse orthologus. Annotation of tumor-infiltrating T cell types was done using ProjectTIL code[6]. Enrichment of GO terms was assessed using the EnrichGO function in the ClusterProfiler package [7,8].

***Virus production for human T cells***

For co-expression of GH1-1, CDT-1, and CD19 CAR, the following viral vectors were used: pYC_cdt1_mcherry, MSCV_gh1-1_GFP, and pYC_CAR_eGFR. To begin viral production, 106 GP2-293 packaging cells (Cell Biolabs) were plated into 6-well plates in 2 mL DMEM supplemented with 10% heat-inactivated fetal bovine serum (FBS) (antibiotics were omitted from this media to increase transfection efficiency and reduce toxicity) and incubated at 37 °C, 5% CO2. 24 h later, GP2 cells were transfected with GH1-1, CDT-1, or CD19 CAR vectors with Lipofectamine 3000 (Thermo Fisher Scientific) according to manufacturer instructions. The next morning, the media was aspirated and replaced with 3 mL fresh media containing 1.5X ViralBoost (Cat# VB100; ALSTEM). 24 h later, the media was harvested and spun at 500 × g for 5 min (to remove cellular contamination). To generate media for co-transduction with three viruses, supernatants from GH1-1 (GFP), CDT-1 (mCherry), and CD19 CAR (hEGFR) transfected cells were combined. For control cells expressing CD19 CAR, but not GH1-1 or CDT-1, supernatants from CAR (hEGFR) transfected cells was used alone. Viral supernatants were loaded into Amicon Ultra-Centrifugation Filters with a 100k Dalton molecular weight cut-off (Millipore Sigma) before spinning at 1,000 × g for 20 min to concentrate. The ~200 µL of viral media generated from concentration was then brought up to 500 µL using complete T cell media containing 50 U/mL of human recombinant interleukin-2 (hrIL-2, BRB Preclinical Repository, National Cancer Institute, NIH). 12-well non-tissue culture treated plates were coated with 50 µg/mL RetroNectin (Takara Bio). The viral supernatants were then added to these wells and centrifuged at 1,000 × g for 90 min. To increase transduction efficiency, viral media was left in the wells before overlaying T cell suspensions (below). To further ensure high levels of transduction efficiency, viral supernatants were always used for transduction on the same day that the supernatants were harvested.

***Human T cell culture, transduction, and expansion***

T cells were isolated from the peripheral blood mononuclear cells of healthy donors using an EasySep immunomagnetic negative selection enrichment kit (Stem Cell Technologies, Cat# 17951). Complete T cell media for our experiments comprised RPMI-1640 supplemented with 10% heat-inactivated FBS, 100 U/mL penicillin/streptomycin, 1 mM sodium pyruvate, 10 mM 4-(2-hydroxyethyl)-1-piperazineethanesulfonic acid (HEPES) buffer, and 55 µM β-mercaptoethanol. Tissue culture-treated 24-well plates were coated with 1 µg/mL anti-CD3 (clone OKT3; Biolegend) overnight at 4 °C then washed. T cells were activated by resuspending at a density of $10^6$ cells/well in complete T cell media additionally containing 2 µg/mL of soluble anti-CD28 (clone CD28.2; Biolegend) and 50 U/mL of hrIL-2. 1 mL was plated onto each well, and then incubated at 37 °C, 5% CO2. After 24 h, T cells were harvested and resuspended in an equal volume of fresh complete T cell media with 50 U/mL of hrIL-2 and 1 mL of T-cell suspension was overlaid onto virus-loaded plates (see Virus production above). T cells were "spinfected" by centrifuging the plates at 1,000 × g at 32 °C for 90 min. Then the plates were incubated in the 37 °C incubator for an additional 90 min. The viral media was then carefully removed, leaving behind the adherent

T cells, before adding back fresh media with 50 U/mL of hrIL-2. Transduced T cells were incubated at 37 °C overnight. The next morning, T cells were harvested from each well and transferred into 15 mL of fresh complete T cell media supplemented with 50 U/mL of hrIL-2 before incubating at 37 °C for another 24 h. Three days post-transduction, T cells were harvested and resuspended in fluorescence associated cell sorting (FACS) buffer (comprising Dulbecco's PBS supplemented with 2% FBS and 1 mM ethylenediaminetetraacetic acid (EDTA)) and stained for CAR transduction with anti-hEGFR PE (clone Hu1; R&D Systems). Triple positive cells (GFP+ mCherry+ PE+) expressing GH1-1, CDT-1, and the CAR receptor were sorted using a Sony SH800S flow cytometric sorter. After sorting, cells were incubated with CD3/CD28 Dynabeads (Thermo, catalog number 11131D, 1:1 ratio). Expansion involved passaging of T cells into fresh complete T cell media with 50 U/mL of hrIL-2 daily. Experiments were performed approximately 12-14 days after T-cell activation.

***Human CAR-T cell functional assays (proliferation, viability, killing, cytokine production)***

T cells were activated and transduced (either with the CAR construct alone or with the CAR construct plus CDT-1 and GH1-1) and expanded with IL-2 as above. To assess their function, T cells were then washed and incubated in media conditions that exposed them to IL-2 plus specific metabolic constraints (e.g., low glucose, with or without cellobiose) overnight. Then they were recounted and adjusted for equal numbers. Then various numbers of T cells were mixed with a 1:1 mixture of Raji cells either lacking CD19 (which lack GFP) or expressing CD19 (which also express GFP). The exact ratio of Raji cells was measured experimentally. The various ratios of T-to-Raji cells were cultured overnight in the same metabolic conditions. Killing of the Raji-CD19 cells (which represented specific killing) versus the Raji cells lacking CD19 (which represented non-specific background death of the cells during culture) were analyzed by flow cytometry, first gating on human CD20 and then separating the two populations by GFP expression. Separately, to assess cytokine production, transduced T cells were co-cultured with only Raji-CD19 cells overnight in metabolic conditions, and the resulting supernatants were assayed for IFNγ and TNF production by cytokine bead array (BD). Separately, to assess viability and proliferation, transduced T cells were labeled with CellTrace Violet (CTV, as per manufacturer's directions) and co-cultured with only Raji-CD19 cells for 48 h in metabolic conditions. T cells were assayed by flow cytometry for viability with a live-dead stain, and proliferation gating on the T cells by human CD3 staining and then measuring geometric mean of CTV. Rescue of proliferation was reported as CTV normalized to a low glucose condition, where proliferation was impaired.

***Raji-CD19 tumor model***

For tumor experiments, we used adult female NCG mice as recipients. Nair was placed on the lower right abdominal area to remove hair. Raji cells were suspended in HBSS and injected subcutaneously. The tumors were allowed to grow until day 10, then mice were reassorted into two groups such that the average size of the tumors was the same in both groups. The mice then underwent i.v. injection of human CAR-CG-T cells. Starting on the day of T-cell transfer, the mice were injected three-times daily with cellobiose or PBS as above. Human anti-PD1 (pembrolizumab) 250 μg was injected i.p. on days 12, 15, and 18. Tumor sizes were followed by calipers, with volume calculated as above. The physical health and behavior of mice were monitored daily as above. Mice were euthanized, tumors removed and suspended in formalin and submitted to the core facility for immunohistochemistry staining for human CD3, human CD19, and human Ki67.

***Microbiome***

NSC mice were subcutaneously injected with Raji-hCD19 cells tumors as above. Tumors were allowed to grow for 10 days, then the mice were re-assorted so that the average tumor size was the same between two groups. The mice were moved to singly housed caging. No T cells were transferred. We injected mice repeatedly with either cellobiose or PBS as above. After 10 days, scat pellets were collected in sterile containers. DNA was prepared using the Mouse QIAamp DNA stool mini kits. We used Zymo Research (Irvine, CA) services for sequencing bacterial and fungal genomes. Bacterial 16S ribosomal RNA gene targeted sequencing was performed using the Quick-16S NGS Library Prep Kit. In most cases, the bacterial 16S primers amplified the V3-V4 region of the 16S rRNA gene. These primers were customdesigned by Zymo Research to provide the best coverage of the 16S gene while maintaining high sensitivity. Fungal ITS gene targeted sequencing was performed using the Quick-16S NGS Library Prep Kit with custom ITS2 primers substituted for 16S primers. The sequencing library was prepared using a custom library preparation process in which PCR reactions were performed in real-time PCR machines to control cycles and therefore limit PCR chimera formation. The final PCR products were quantified with qPCR fluorescence readings and pooled together based on equal molarity. The final pooled library was cleaned with the Select-a-Size DNA Clean & Concentrator (Zymo Research, Irvine, CA), then quantified with TapeStation. The final library was sequenced on Illumina Nextseq with a P1 reagent kit (600 cycles). The sequencing was performed with 30% PhiX spike-in. Analyses (e.g., heatmaps) were performed using proprietary scripts at Zymo Research and raw data replotted and analyzed in R. P values for diversity indices were calculated using ANOVA (diversity indices versus variates of mice and treatments).

### Supplemental Figures and Legends

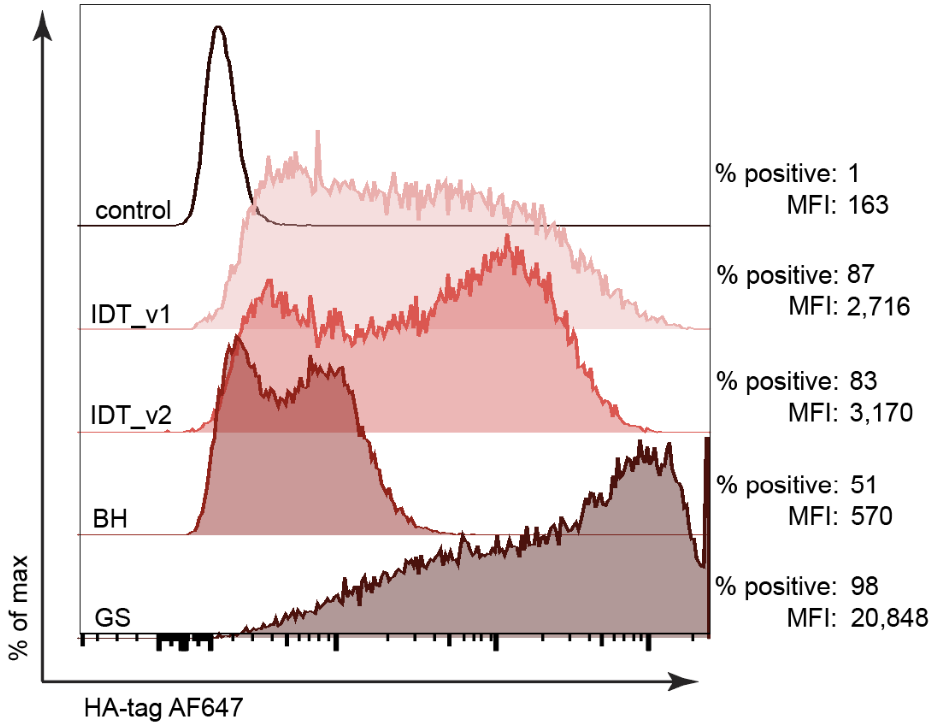


**Fig. S1. Codon optimization approaches for CDT-1 expression.** To generate codon sequences for mammalian expression of CDT-1, we employed publicly available codon optimization algorithms from Integrated DNA Technologies (IDT), Blue Heron Biotech (BH), and GenScript (GS). The sequences were cloned into an MSCV expression vector and transfected into Platinum-E (HEK 293T) cells. After 48 h, the cells were harvested, fixed, permeabilized, and immunostained for the addended HA-tag. The sequence generated by the GenScript codon optimization algorithm led to the highest expression of CDT-1.

sdfsdaf

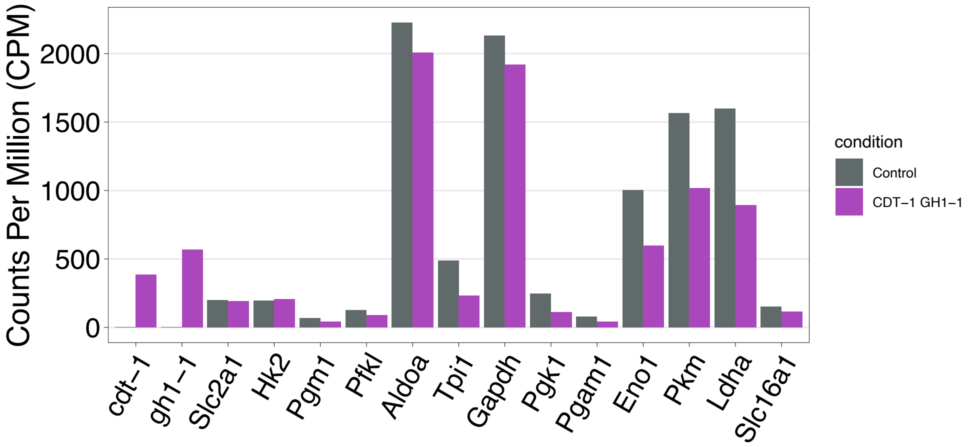


**Fig. S2. Bulk RNAseq showing no impact on metabolic pathways by introduction of CG transgenes.** Transcription of CG-transduced T cells versus control transduced T cells. Transcripts of the two transgenes cdt-1 and gh1-1 as well as the fluorophore mCherry were highly enriched. Shown are selected metabolic genes, with no statistically significant differences. Bulk RNAseq revealed no statistically significant enrichment of any Reactome pathways.

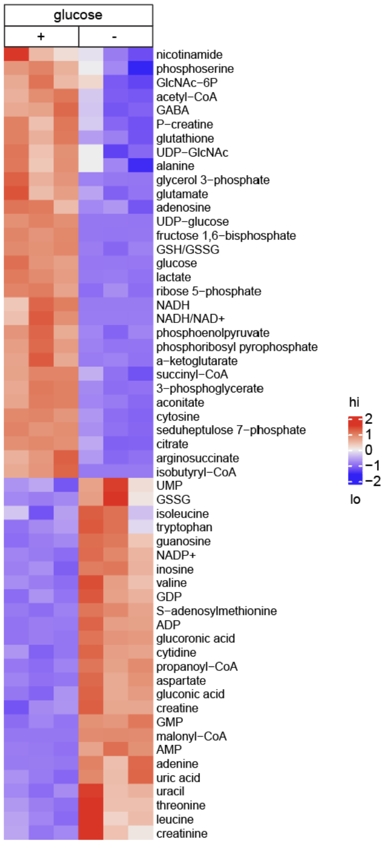


**Fig. S3. Glucose deprivation leads to profound metabolic disturbances in T cells.** Activated T cells were cultured with and without glucose for 16 h and analyzed by LC-MS. T cells deprived of glucose showed depletion of key nutrients in the glycolytic, pentose phosphate, glycosylation and TCA pathways. T cells deprived of glucose accumulated low-energy and oxidative intermediates including AMP and oxidized glutathione (GSSG).

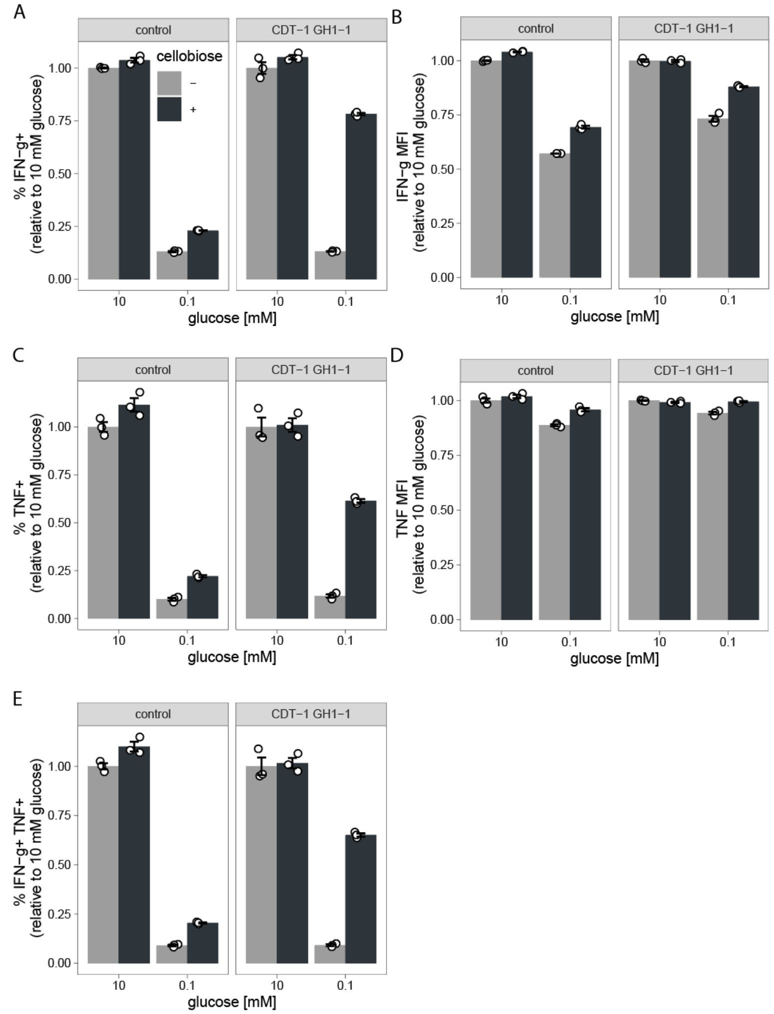


**Fig. S4. Cellobiose rescues intracellular cytokine levels in glucose-deprived CG-T cells.** Control and CG-T cells were incubated for 16 h in the indicated metabolic conditions before stimulation with PMA/ionomycin in the presence of Brefeldin A. The cells were then fixed, permeabilized, and stained to measure the level of intracellular cytokines. The percentages of cells expressing IFNγ and TNF were increased by cellobiose in CG-T cells, but not control cells. Data in this experiment are means of technical replicates (n=3) and representative of at least two separate experiments.

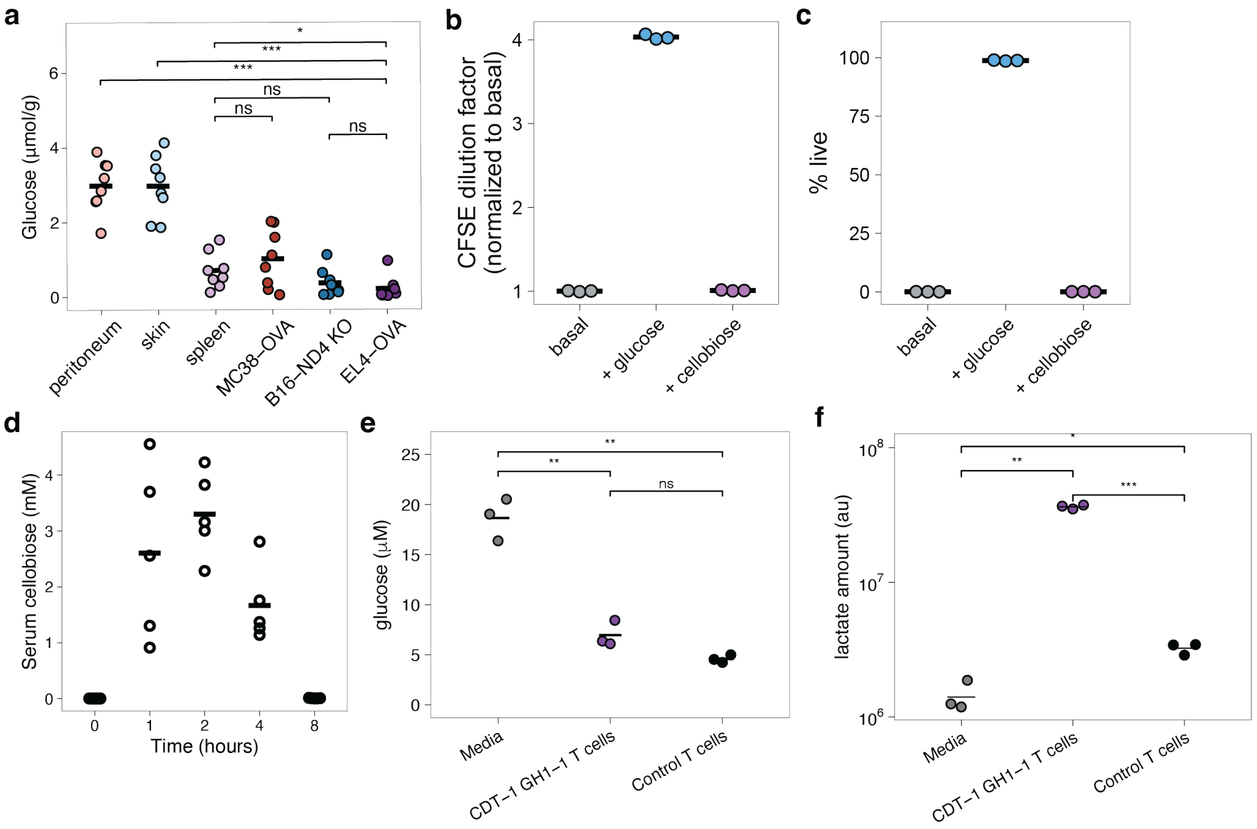


**Fig. S5. Tumors cannot metabolize cellobiose and pharmacokinetics; T cells cannot provide glucose to tumor cells. a)** Tumors and healthy tissues measured for glucose levels by LC-MS. **b)** Proliferation of EL4-OVA tumors cultured with media deprived of glucose (basal), with glucose and with cellobiose. More CFSE dilution indicates more proliferation. **c)** Survival of EL4-OVA tumors cultured with media deprived of glucose (basal), with glucose and with cellobiose as measured by flow cytometry. **d)** Serum concentration of cellobiose at various timepoints after i.p. injection of 85 mg of cellobiose (n=5). Statistical significance in (**a**) was assessed using an unpaired t-test in (*$p < 0.05$; **$p < 0.01$; ***$p < 0.001$; NS > 0.05). **e** and **f)** Mouse CD8+ T cells were transduced with either control viruses or CDT-1 and GH1-T, cultured, and sorted for purity before incubating in 5 mM cellobiose medium overnight. LC-MS was run on the media to assess glucose and lactate. 5 mM cellobiose media is contaminated with a measurable level of glucose. Results show three parallel, independent T cell transductions performed at the same time (but not pipetting replicates), and bar indicates average. Glucose measurements were calibrated using $^{13}$C-glucose spike-in controls. Lactate measurements are raw counts from MS.

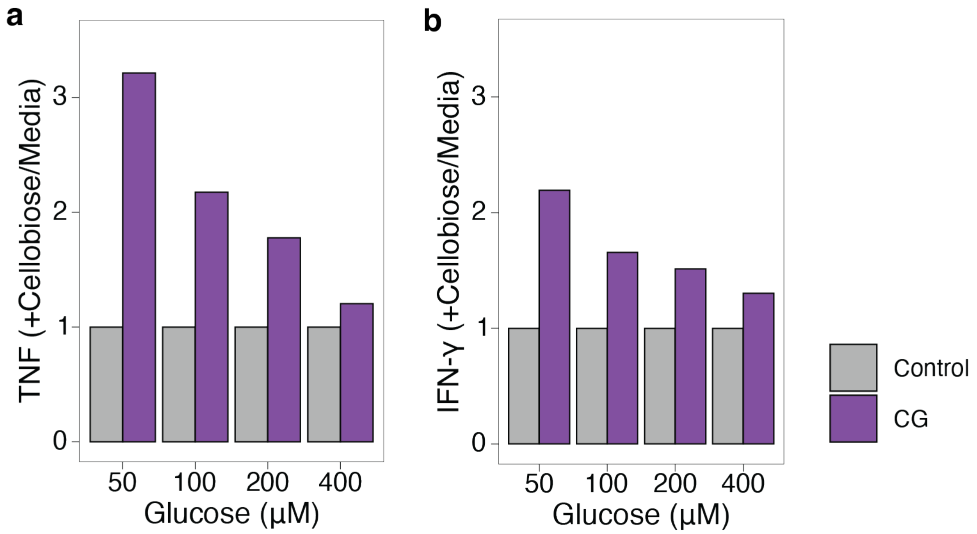


**Fig. S6. CAR-CG-T cells show an advantage in low glucose.** CAR-CG and CAR-only transduced T cells cultured in 0.1 mM glucose overnight, then cultured with target cells overnight in various glucose and cellobiose concentrations as indicated. Cytokine production in the supernatant was measured by cytokine bead array. To highlight the accentuation by cellobiose in CAR-CG-T cells over T cells transduced with the CAR alone, we divided the cytokine amount (in pg/mL) in the media+cellobiose condition by the control media condition. Then we normalized this ratio for the CAR-CG-T cells to the ratio for the CAR alone T cells. In all glucose concentrations, CAR-CG-T cells showed more enhancement in the presence of cellobiose than did the CAR alone T cells.

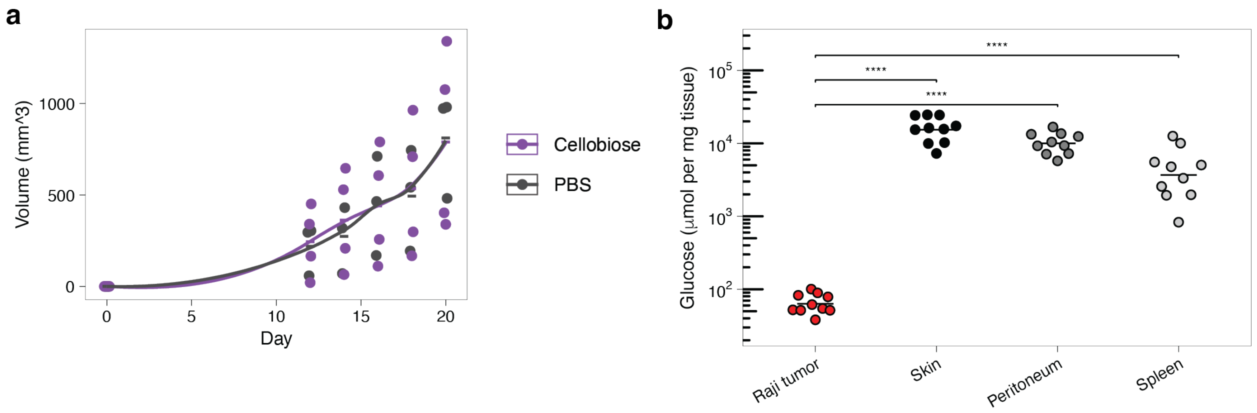


**Fig. S7. Raji tumors are not benefited by cellobiose and thrive in a low glucose environment in vivo. b)** Raji tumors were subcutaneously implanted in mice. Once tumors were established, mice were reassorted into two groups with equal average tumor size. One group received cellobiose injections and the other an equal volume of saline. No engineered T cells were given. Tumor sizes were measured by calipers. **a)** Subcutaneously implanted Raji tumors were dissected and tissues measured for glucose levels by LC-MS, as compared to healthy tissues. Skin was adjacent to the subcutaneous tumors.

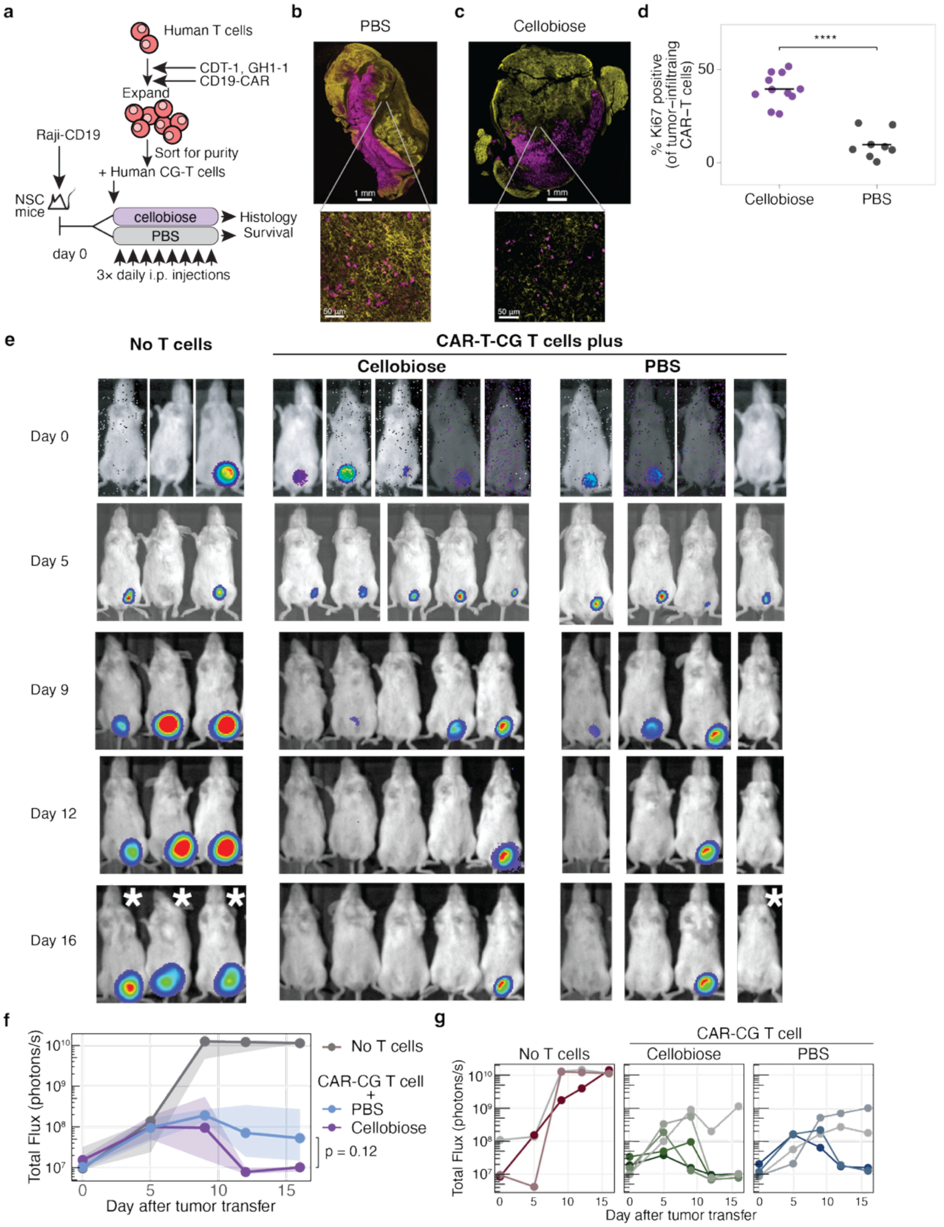


**Fig. S8. Proliferation of human CAR-T cells in tumor. a)** Human CD8 T cells are transduced with the CDT-1 and GH1-1 genes and CD19-CAR, then transferred into immunodeficient mice bearing Raji-CD19 luc tumors. Mice are treated with cellobiose or PBS. **b** and **c)** Tumor cross sections were immunostained for human CD19 (Raji cells, false-colored yellow), human CD3 (CAR-T cells, false-colored magenta), and Ki67. Zoomed sections show individual tumor-infiltrating T cells that were adjacent to sheets of T cells and that were studied for Ki67 expression. **d)** Ki67 positive T cells from tumor infiltrating regions. Each dot represents the TILs captured in non-overlapping regions of approximately 0.6 $mm^2$ in the zone of ~0.5 mm from the T cell-tumor boundary in a single tumor, and thus the individual regions are non-independent. **e)** luciferase imaging of the Raji-CD19 luc tumors at various days after implantation. Mice are grouped by treatments. White asterisk indicates imaging for 10 s (and thus the luciferase image is less intense) whereas the rest of the images were collected for 30 s. **f)** Photon flux from the tumors over time is shown for each group with median values plotted as circles, and first and third quartiles as ribbons. Wilcox p-value shown. **g)** Photon flux from the tumors over time is shown for the individual mice in each group.

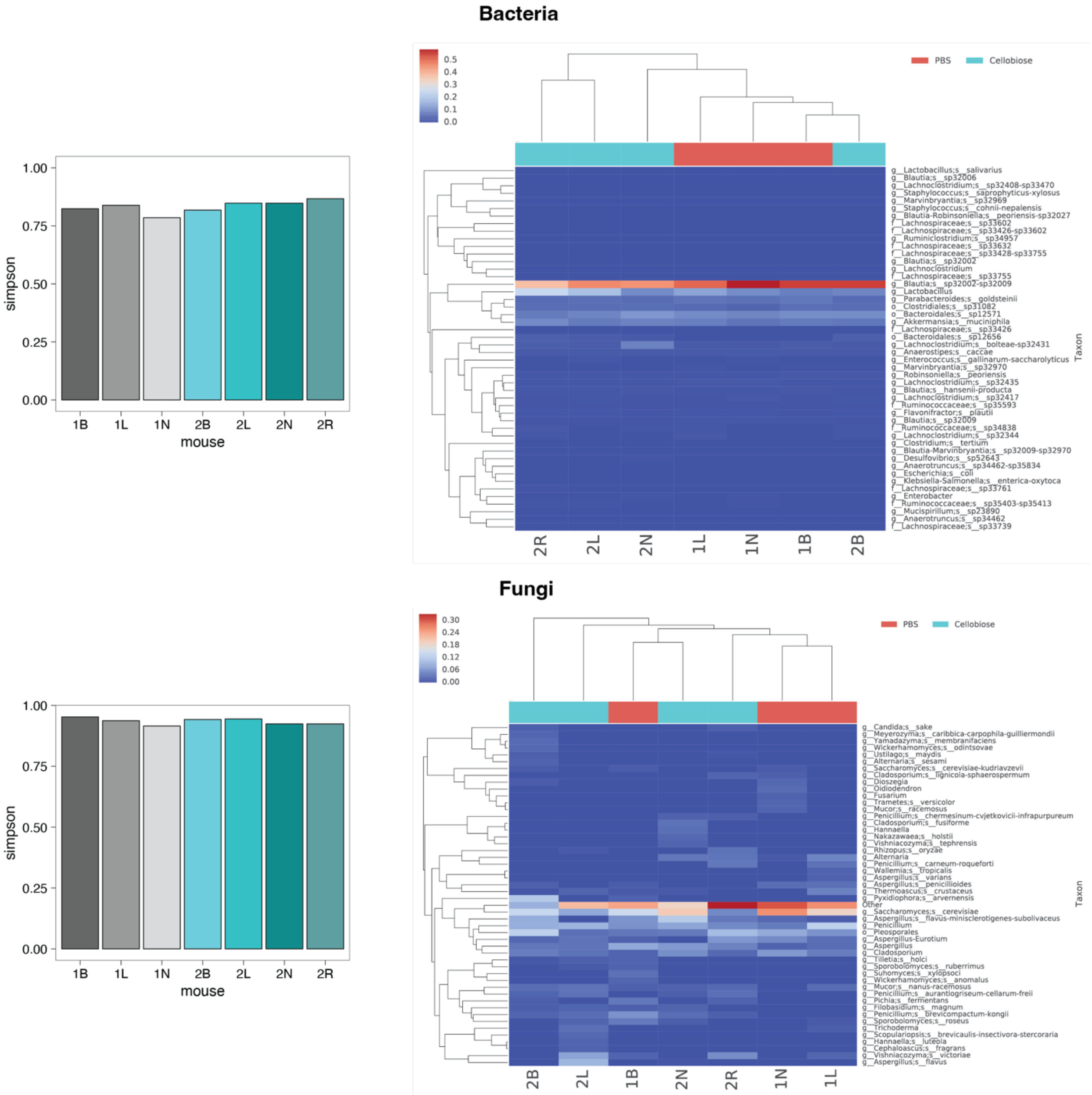


**Fig. S9. Microbiome after cellobiose treatment.** Gut microbiome of bacteria and fungi as assessed by 16S and ITS sequencing from fecal pellets from mice treated with cellobiose or PBS for 10 days. Simpson diversity index is shown on the left. Cyan bars represent individual mice treated with cellobiose, gray are control mice treated with PBS. Clustered heatmaps are shown on the right. The identity of the mice at the top of each heatmap shows PBS treatment (coral) or cellobiose (cyan).
